# Lack of *Tgfbr1* and *Acvr1b* synergistically stimulates myofibre hypertrophy and accelerates muscle regeneration

**DOI:** 10.1101/2021.03.03.433740

**Authors:** M.M.G. Hillege, A. Shi, R.A. Galli, G. Wu, P. Bertolino, W.M.H. Hoogaars, R.T. Jaspers

**Author notes:** Contributed equally to this manuscript (M.M.G. Hillege, A. Shi).

## Abstract

In skeletal muscle, transforming growth factor-β (TGF-β) family growth factors, TGF-β1 and myostatin, are involved in atrophy and muscle wasting disorders. Simultaneous interference with their signalling pathways may improve muscle function, however little is known about their individual and combined receptor signalling. Here we show that inhibition of TGF-β signalling by simultaneous muscle-specific knockout of TGF-β type I receptors *Tgfbr1* and *Acvr1b* simultaneously, induces substantial hypertrophy, while such effect does not occur by single receptor knockout. Hypertrophy is induced by increased phosphorylation of Akt and p70S6K and reduced E3 ligases expression, while myonuclear number remains unaltered. Combined knockout of both TGF-β type I receptors increases the number of satellite cells, macrophages and improves regeneration post cardiotoxin-induced injury by stimulating myogenic differentiation. Extra cellular matrix gene expression is exclusively elevated in muscle with combined receptor knockout. *Tgfbr1* and *Acvr1b* are synergistically involved in regulation of myofibre size, regeneration and collagen deposition.

## Introduction

Muscle wasting disorders, such as muscular dystrophies, cancer cachexia and sarcopenia are characterised by reduced muscle mass, impaired regeneration and fibrosis, which results in progressive muscle weakness. The transforming growth factor β (TGF-β) superfamily members TGF-β1, myostatin and activin A are involved in various processes within muscle tissue and overexpression of these proteins contributes to muscle wasting pathologies (Bernasconi et al., 1995; Carlson, Hsu, & Conboy, 2008; J. L. Chen et al., 2014; Costelli et al., 2008; Leger, Derave, De Bock, Hespel, & Russell, 2008; Tanaka, Miyazaki, Takeda, & Takeo, 1993).

TGF-β signalling negatively affects muscle growth by both affecting satellite cells (SCs) and myofibres. TGF-β1, myostatin and activin A inhibit myoblast differentiation (Langley et al., 2002; Liu, Black, & Derynck, 2001; Trendelenburg, Meyer, Jacobi, Feige, & Glass, 2012). Inhibition of myostatin and activin A synergistically results in muscle hypertrophy (Amirouche et al., 2009; J. L. Chen et al., 2017; J. L. Chen et al., 2014; Latres et al., 2017; McFarlane et al., 2006; Zimmers et al., 2002). These effects are at least partly independent of SCs (Lee et al., 2012). In addition, TGF-β1 overexpression *in vivo* may also cause muscle atrophy (Mendias et al., 2012; Narola, Pandey, Glick, & Chen, 2013).

Transient TGF-β1 expression may play an essential role during muscle regeneration. TGF-β1 is expressed by inflammatory cells, such as macrophages, monocytes and neutrophils as well as by fibroblasts after acute injury (Assoian et al., 1987; Grotendorst, Smale, & Pencev, 1989; Lawrence, Pircher, Kryceve-Martinerie, & Jullien, 1984; Zimowska et al., 2009). During muscle regeneration, TGF-β1 is involved in the regulation of the immune response and plays an important role in rebuilding extracellular matrix (ECM) (Gillies & Lieber, 2011; Kehrl et al., 1986; Reibman et al., 1991; Tsunawaki, Sporn, Ding, & Nathan, 1988; Wahl et al., 1987; Wiseman, Polverini, Kamp, & Leibovich, 1988). However, chronic increased expression of TGF-β is known to contribute to muscle fibrosis (Y. Li et al., 2004). Furthermore, myostatin and activin A have also been suggested to induce substantial skeletal muscle fibrosis (J. L. Chen et al., 2014; Z. B. Li, Kollias, & Wagner, 2008). Thus, inhibition of TGF-β signalling in the myofibre may substantially reduce connective tissue deposition.

In animal models, such as murine X-linked muscular dystrophy (mdx) mice, a Duchenne muscular dystrophy (DMD) mouse model, cancer cachexia mouse models or aged mice, inhibiting signalling of one or more of these ligands had beneficial effects, such as reduction in fibrosis and maintenance of muscle mass (Andreetta et al., 2006; J. L. Chen et al., 2017; Greco et al., 2015; Latres et al., 2015; K. T. Murphy et al., 2011). Lack of either TGF-β or myostatin has been suggested to improve regeneration after acute injury (Accornero et al., 2014; McCroskery et al., 2005). Therefore, inhibiting these growth factors may be a promising therapeutic strategy to alleviate muscle wasting pathologies.

However, interference with signalling of TGF-β family members may be complicated. TGF-β family members regulate various cellular processes throughout the body, thus systemic inhibition of these growth factors may have severe consequences. Furthermore, due to overlap in function of these ligands, inhibition of a single ligand is likely not effective.

Simultaneous inhibition of TGF-β1, myostatin and activin A through interference with their downstream receptors may be an effective approach. The TGF-β family consists of at least 33 cytokines that can roughly be divided into the TGF-β/myostatin/activins subgroup and the growth differentiation factor/bone morphogenetic protein (BMP) group that often have opposing effects. These cytokines regulate gene expression via specific binding to distinct type II and type I receptors. TGF-β1 mainly signals via the type II receptor, TGF-β receptor type-2 (TGFBR2), and via the type I receptor, TGF-β receptor type-1 (TGFBR1 or ALK5). Myostatin signals via the type II receptor, activin receptor type-2B (ACVR2B), and activin A signals via type II receptors, activin receptor type-2A (ACVR2A) and ACVR2B. Activin A signals via type I receptor, activin receptor type-1B (ACVR1B or ALK4). Myostatin has been shown to signal via both TGFBR1 and ACVR1B in various cell types (Kemaladewi et al., 2012; Rebbapragada, Benchabane, Wrana, Celeste, & Attisano, 2003; ten Dijke et al., 1994).

Interference with myostatin/activin A signalling by blocking their type II receptors ACVR2A/B may not be an appropriate strategy, since these receptors are also involved in BMP signalling, which stimulates muscle hypertrophy whereas myostatin/activin A signalling is associated with atrophy (Sartori, Gregorevic, & Sandri, 2014; Tsuchida, Nakatani, Uezumi, Murakami, & Cui, 2008). Moreover, interference with signalling via these receptors may cause severe side effects, such as nose and gum bleeds, as has been shown in in DMD boys treated with soluble ACVR2B (Campbell et al., 2017). Inhibition of type I receptors TGFBR1 and ACVR1B may provide a more specific and effective approach to alleviate muscle wasting pathologies (Sartori et al., 2014; Tsuchida et al., 2008). However, very little is known about the role of these receptors in the regulation of skeletal muscle mass and regeneration.

The aim of this study was to obtain insight in how myofibre specific knockout of type I receptors *Tgfbr1* and *Acvr1b* affected muscle size as well as early muscle regeneration, inflammation and collagen deposition in both intact and injured muscle. We hypothesised that individual knockout of these TGF-β type I receptors would have marginal effects. Moreover, simultaneous inhibition of these type I receptors would substantially increase muscle size and enhanced early myofibre regeneration, while attenuating fibrosis.

## Results

### *Acvr1b* and *Tgfbr1* expression was successfully reduced after tamoxifen treatment

The aim of this study was to investigate effects of mature myofibre specific knockout of *Tgfbr1* and *Acvr1b* on muscle morphology as well as early muscle regeneration, inflammation and collagen deposition in both uninjured muscle tissue and after acute cardiotoxin (CTX) injury. For this purpose, the HSA-MCM mouse line (McCarthy, Srikuea, Kirby, Peterson, & Esser, 2012b), that expresses tamoxifen (TMX) inducible Cre (MerCreMer) under a human α-skeletal actin (HSA) promotor was cross bred with the conditional knockout Acvr1b^fl/fl^ (Ripoche et al., 2013) and Tgfbr1^fl/fl^ (Larsson et al., 2001) mouse lines to obtain mouse lines HSA-MCM:Acvr1b^fl/fl^, HSA-MCM:Tgfbr1^fl/fl^ and HSA-MCM:Acvr1b^fl/fl^:Tgfbr1^fl/fl^ (further referred to as Acvr1b^fl/fl^, Tgfbr1^fl/fl^ and Acvr1b^fl/fl^:Tgfbr1^fl/fl^). Receptors were deleted when mice were 6 weeks old (Figure 1A and B).

**Figure 1.**
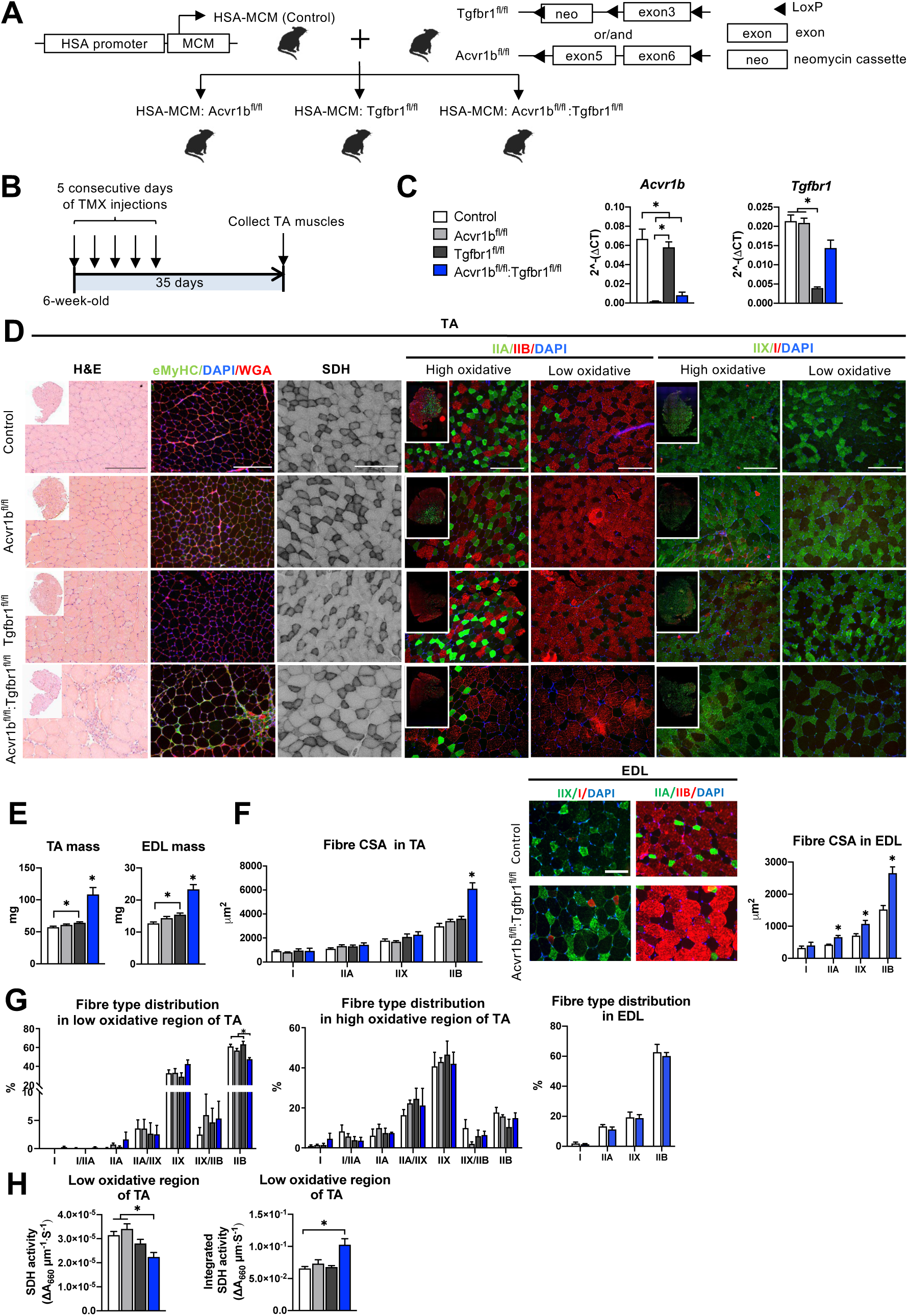
Simultaneous knockout of both *Acvr1b* and *Tgfbr1* caused muscle hypertrophy. (A) Scheme showing cross-breeding of HSA-MCM mouse line with conditional knockout mouse lines Acvr1b^fl/fl^ and Tgfbr1^fl/fl^. LoxP sites are indicated by black arrows. A loxP-flanked neomycin (neo) cassette is inserted upstream of exon3 of Acvr1b genome. (B) Scheme demonstrating receptor knockout induced by tamoxifen (TMX) injection for consecutive 5 days. (C) Relative mRNA expression of *Acvr1b* and *Tgfbr1* in TA muscles of experimental groups. (D) Histology stainings of TA muscles 35 day after first TMX injection. H&E staining and immunofluorescent staining of eMyHC (green) of TA showed regenerative regions containing eMyHC^+^ myofibres with central nuclei (DAPI, blue) in Acvr1b^fl/fl^:Tgfbr1^fl/fl^ mice, wheat glucose agglutinin (WGA, red) was used to visualise cell membranes and ECM. Acvr1b^fl/fl^:Tgfbr1^fl/fl^ mice showed lower staining intensity for SDH in low oxidative region of TA. MyHCs staining demonstrated type IIA (green), IIB (red), IIX (green) and I (red) myofibres in low and high oxidative regions of TA. Scale bars=250 μm. (E) TA and EDL muscle masses and myofibre cross-sectional areas (CSAs) were increased in Acvr1b^fl/fl^:Tgfbr1^fl/fl^ mice. (F) In TA, specifically CSA of type IIB myofibres was increased in Acvr1b^fl/fl^:Tgfbr1^fl/fl^ animals, while in EDL CSA of all type II myofibres was increased. Myofibre types were stained in EDL. (G) Percentage of type IIB in low oxidative region of TA was reduced. No differences were observed in myofibre distribution in high oxidative region of TA or EDL. (H) SDH activity (absorbance units (ΔA_660_) per micrometer section thickness per second of incubation time (ΔA_660_·μm^-1^·s^-1^)) was decreased, while the integrated SDH activity SDH activity multiplied by CSA (ΔA_660_·μm·s^-1^) increased in low oxidative region of TA of Acvr1b^fl/fl^:Tgfbr1^fl/fl^ animals. N=5-8 mice. Results are presented as mean + SEM. *: *P* < 0.05. Significant difference between individual groups is indicated by lines with a *. Single * indicates significant difference compared to all other groups.

Expression levels of *Acvr1b* and *Tgfbr1* mRNA showed successful knockout as *Acvr1b* mRNA levels in tibialis anterior (TA) muscles were reduced in Acvr1b^fl/fl^ animals by 97% and in Acvr1b^fl/fl^:Tgfbr1^fl/fl^ animals by 88%. *Tgfbr1* expression levels in TA muscles were reduced in Tgfbr1^fl/fl^ animals by 82% Unexpectedly, *Tgfbr1* expression levels in TA muscle of Acvr1b^fl/fl^:Tgfbr1^fl/fl^ animals were not significantly reduced compared to those of control animals (Figure 1C). Note, however, that lack of significantly reduced *Tgfbr1* expression is likely a consequence of high *Tgfbr1* expression by other cell types present within the muscle, rather than unsuccessful knockdown. This issue is addressed below in more detail (see Figure 3). *Acvr1b* expression levels did not affect *Tgfbr1* expression levels and vice versa.

### Simultaneous knockout of *Acvr1b* and *Tgfbr1* resulted in type IIB myofibre hypertrophy and has modest effects on myofibre type distribution

TGFBR1 and ACVR1B ligands are well known for their regulatory effects on muscle mass. Here, TA mass of Acvr1b^fl/fl^:Tgfbr1^fl/fl^ mice (108.4 ± 11.0 mg) was nearly doubled compared to that of control animals (57.2 ± 1.5 mg). TA mass of Tgfbr1^fl/fl^ mice (64.2 ± 1.4 mg) was also increased, however to a much lower extend. TA mass of Acvr1b^fl/fl^ mice (60.8 ± 1.5 mg) did not differ from that of controls (Figure 1E). To test whether the observed effects in TA also apply to other muscles, extensor digitorum longus muscle (EDL) mass was determined. Similar to TA muscle, EDL mass of Acvr1b^fl/fl^:Tgfbr1^fl/fl^ (23.3 ± 1.5 mg) and Tgfbr1^flfl^ mice (15.4 ± 0.5 mg) was increased by 1.8-fold and 1.3-fold, compared to that of control mice (12.7 ± 0.4 mg), respectively, while EDL mass of Acvr1b^fl/fl^ mice (14.4 ± 0.5 mg) did not differ from that of control mice (Figure 1E). Note that in TA, specifically the cross-sectional area (CSA) of type IIB myofibres of Acvr1b^fl/fl^:Tgfbr1^fl/fl^ mice was twofold larger compared to that of control animals (Figure 1F), indicating that simultaneous knockout of both *Acvr1b* and *Tgfbr1* synergistically causes myofibre hypertrophy in type IIB myofibres. In contrast to observations in TA, CSA of type IIA and type IIX myofibres in EDL muscle of Acvr1b^fl/fl^:Tgfbr1^fl/fl^ mice were increased by 1.6-fold and 1.5-fold compared to those of control mice, respectively. However, similar to TA muscle, CSA of type IIB myofibres of Acvr1b^fl/fl^:Tgfbr1^fl/fl^ mice was increased most substantially compared to that of control mice (1.7-fold) (Figure 1F).

These results indicate that simultaneous knockout of *Acvr1b* and *Tgfbr1* in mature mouse myofibre synergistically causes muscle hypertrophy of mostly type IIB myofibres, whereas individual knockout has little effect on muscle mass or myofibre CSA.

Observed effects on myofibre CSA may indicate alterations in myofibre metabolism as well as in myofibre type distribution. Therefore, myofibre type distribution was determined in both the high and low oxidative region of the TA (Figure 1D and G). In the high oxidative region no significant differences were observed. However, in the low oxidative region of Acvr1b^fl/fl^:Tgfbr1^fl/fl^ animals the percentage of type IIB myofibres was lower (47 ± 2%) compared to both control (61 ± 2%) and Tgfbr1^flfl^ animals (63 ± 3%) (Figure 1G), which indicates a shift towards an oxidative phenotype in these muscles. In contrast, no differences in EDL myofibre type distribution were observed between Acvr1b^fl/fl^:Tgfbr1^fl/fl^ and control mice (Figure 1F and G).

Taken together, lack of both *Acvr1b* and *Tgfbr1* reduces the percentage of type IIB myofibres in TA muscle, but has only modest effects on myofibre type distribution.

### Type IIB myofibre hypertrophy resulted in reduced SDH activity

In skeletal muscle, myofibre size and oxidative capacity are inversely related, indicating that metabolism implies a size constraint (W. van der laarse, Tombe, Groot, & Diegenbach, 1997; van Wessel, de Haan, van der Laarse, & Jaspers, 2010). To test whether the excessive hypertrophy within the low oxidative region of the TA muscles was accompanied by a reduction in oxidative metabolism, succinate dehydrogenase (SDH) activity and integrated SDH activity were determined. In Acvr1b^fl/fl^:Tgfbr1^fl/fl^ mice SDH activity was decreased by 30% compared to that in Acvr1b^fl/fl^ and control animals. However, the integrated SDH activity (total oxidative capacity of myofibres) in Acvr1b^fl/fl^:Tgfbr1^fl/fl^ mice was increased by 60% compared to that in control animals. This suggests that while locally the total oxidative capacity in the low oxidative region of TA muscle of Acvr1b^fl/fl^:Tgfbr1^fl/fl^ animals may be reduced, the oxidative capacity per myofibre in the low oxidative region of TA muscle of these animals was substantially increased (Figure 1H).

### Myofibres with central nuclei and increased number of SCs were observed in TA muscle of Acvr1b^fl/fl^:Tgfbr1^fl/fl^ mice

Hematoxylin & Eosin (H&E) staining and embryonic myosin heavy chain (eMyHC) staining showed within TA of uninjured Acvr1b^fl/fl^:Tgfbr1^fl/fl^ animals, regions with small myofibres with centrally located nuclei, indicating injured myofibres. These myofibres were eMyHC^+^ and surrounded by many other cells, likely a combination of SCs, fibroblasts and immune cells (Figure 1D). These regions with regenerating myofibres were mainly present in the low oxidative region of the TA and comprised on average 2.95% of the muscle CSA. These regions were almost never observed in TA of other animals (<0.2%) (Figure 2A). Similar regions with myofibres containing centrally located myonuclei were also observed in EDL (1.45% of the muscle CSA) of Acvr1b^fl/fl^:Tgfbr1^fl/fl^ animals and not in EDL of control animals (0.08%) (Figure 2A). Together, these data indicate that simultaneous knockout of *Acvr1b* and *Tgfbr1* results in spontaneous damage and regeneration. Spontaneous regeneration requires activation of SCs. In the low oxidative region of TA muscle of Acvr1b^fl/fl^:Tgfbr1^fl/fl^ animals the number of SCs per myofibre in a cross-section was increased compared to that in control animals (Figure 2B).

**Figure 2.**
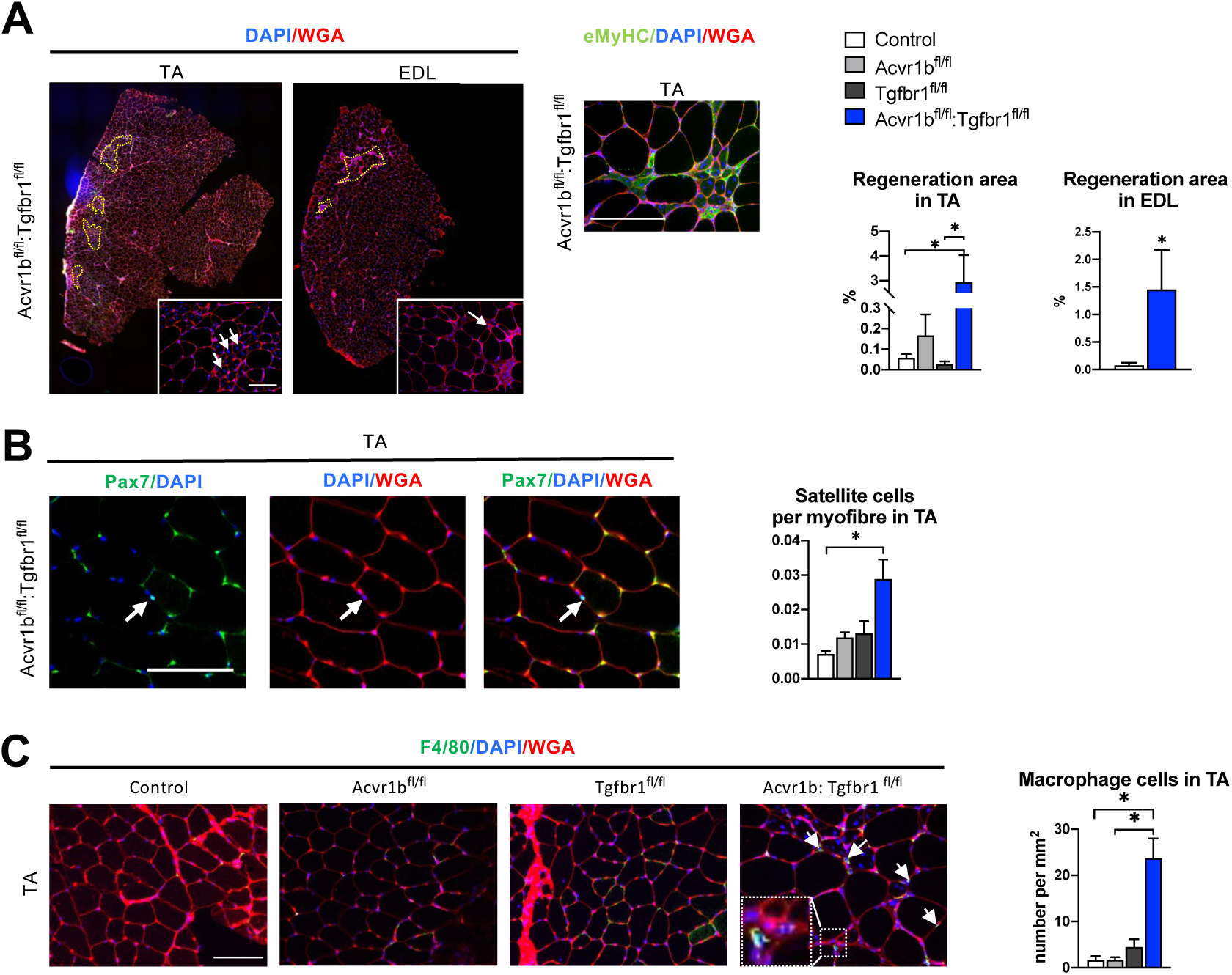
Increased heterogeneity of cell types was found in both TA and EDL of Acvr1b^fl/fl^:Tgfbr1^fl/fl^ animals. (A) Regions with spontaneously regenerating myofibres (circled by yellow dash lines) with central nuclei (indicated by arrows) were particularly present in low oxidative region of TA and EDL of Acvr1b^fl/fl^:Tgfbr1^fl/fl^ animals. (B) Increased number of Pax7^+^ cells per myofibre was found in TA of Acvr1b^fl/fl^:Tgfbr1^fl/fl^ mice. (C) IF staining of F4/80 (green) showed an increased number of macrophages (indicated by arrows) in TA muscle per mm^2^ CSA of Acvr1b^fl/fl^:Tgfbr1^fl/fl^ mice compared to control. Macrophages (image with higher magnification on the left corner) were mainly located around myofibres with central nuclei. Scale bar indicates 100 µm. N=5-8 mice. Results are presented as mean + SEM. *. P < 0.05. Significant difference between individual groups is indicated by lines with a *. Single * indicates significant difference compared to all other groups at the same time point.

We next characterised cells surrounding the spontaneously regenerating regions in TA as being macrophages or fibroblasts. F4/80 staining showed that the number of macrophages per mm^2^ muscle CSA in Acvr1b^fl/fl^:Tgfbr1^fl/fl^ animals was increased by 14-fold (23.7 cells/mm^2^) compared to that in control (1.7 cells/mm^2^) and Acvr1b^fl/fl^ animals (1.7 cells/mm^2^), while in Tgfbr1^fl/fl^ animals (4.5 cells/mm^2^) the number of macrophages per mm^2^ did not differ compared to that in the other three groups (Figure 2C).

Taken together, in TA and EDL muscles that lack both *Acvr1b* and *Tgfbr1* regions with spontaneously regenerating myofibres were observed. These regions are accompanied by an increased number of SCs and macrophages.

### Lack of both *Acvr1b* and *Tgfbr1* in the skeletal myofibre increased *Hgf* expression levels and Akt/p70S6K signalling, while decreasing *Murf-1* expression levels

Next, we aimed to obtain insight in the mechanisms underlying the increase in Acvr1b^fl/fl^:Tgfbr1^fl/fl^ TA mass and myofibre CSA, as well as the observed increase in SC number and regeneration regions in these muscles. First, we determined whether the increase in myofibre size was accompanied by accretion of myonuclei. Counts of myonuclear fragments in muscle cross-sections of IIB myofibres did not differ between control and Acvr1b^fl/fl^:Tgfbr1^fl/fl^ animals. Moreover, in longitudinal sections no difference in length of myonuclei per myofiber was found between groups (Figure 3A), which indicates that the probability to encounter a myonucleus within a cross-section was equal between groups. The excessive hypertrophy of type IIB myofibres of Acvr1b^fl/fl^:Tgfbr1^fl/fl^ mice occurred without accretion of myonuclei and caused a 70% increase in the myonuclear domain (Figure 3A).

**Figure 3.**
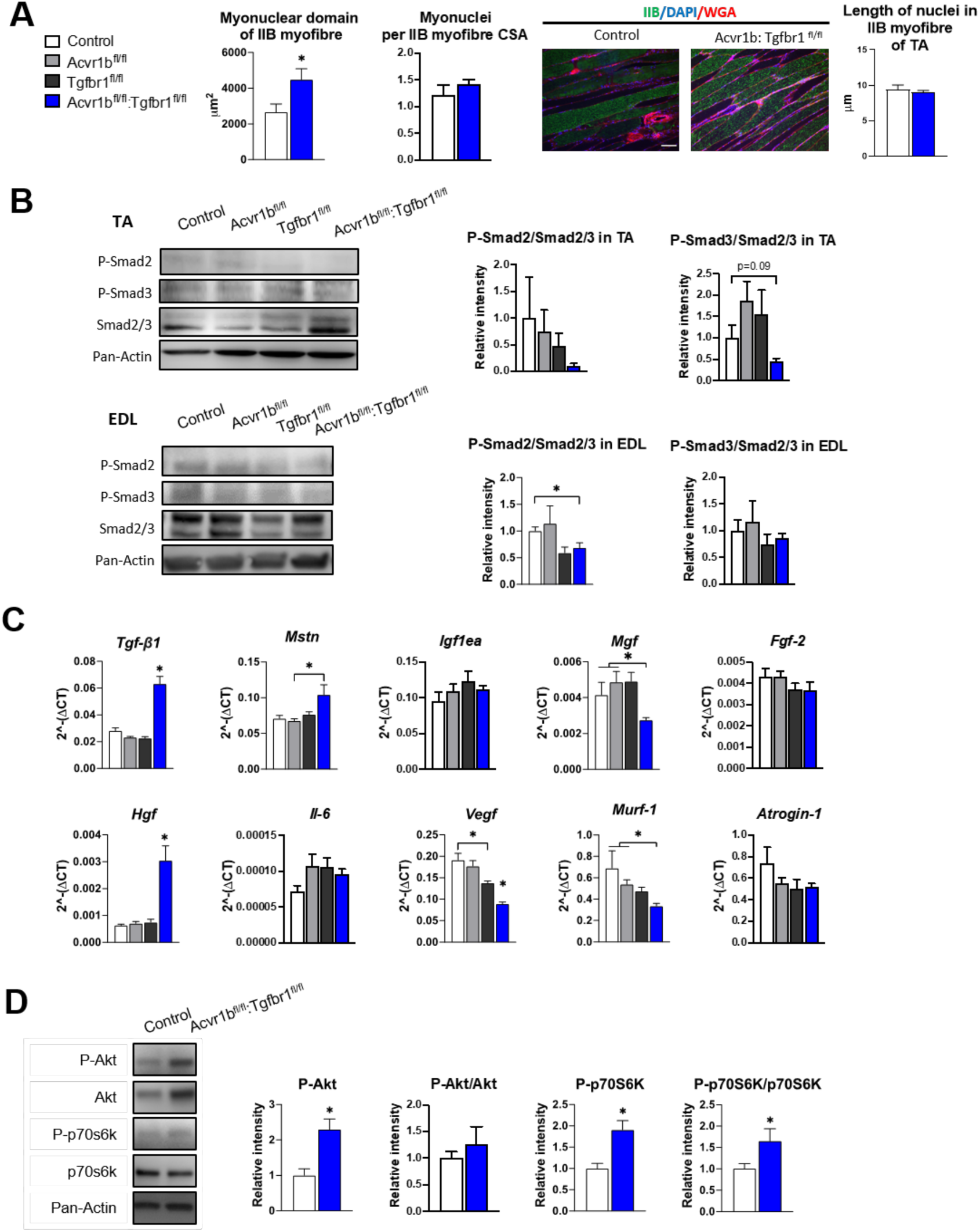
Effects of simultaneous knockout of both *Acvr1b* and *Tgfbr1* on myonuclear number and signalling for protein synthesis as well as degradation. (A) No differences in myonuclear lengths were observed in longitudinal sections of TA IIB myofibres of Acvr1b^fl/fl^:Tgfbr1^fl/fl^ compared to control animals. This indicates that simultaneous knockout of Acvr1b^fl/fl^:Tgfbr1^fl/fl^ did not affect the number of myonuclei per myofibre and that the myonuclear domain (i.e. cross-sectional area/ nuclei (μm^2^)) was almost doubled. Scale bar=100μm. (B) Western blot analysis for Smad2/3 phosphorylation in TA and EDL muscle. (C) Relative gene expression of growth factors in non-injured muscle. (D) Western blot analysis of phosphorylated and total Akt and p70S6K in TA muscles. Results are presented as mean + SEM. N=5-8 mice. *: *P* < 0.05. Significant difference between individual groups is indicated by lines with a *. Single * indicates significant difference compared to all other groups at the same time point.

Phosphorylation of TGF-β type I receptor is known to activate canonical Smad2/3 signalling. Therefore, we examined the effects of TGF-β type I receptor knockout on Smad2/3 phosphorylation in both TA and EDL muscle (Figure 3B). Single knockout did not affect phosphorylated / total protein ratios for Smad2 and Smad3 in muscles of Acvr1b^fl/fl^ or Tgfbr1^fl/fl^ mice, which was in line with the lack of effect on muscle size and phenotype and suggested that at least the presence of one of the two receptors was sufficient to maintain the Smad signalling. With regard to Smad2/3 phosphorylation in TA and EDL of Acvr1b^fl/fl^:Tgfbr1^fl/fl^ mice, a 31% reduction was shown for phosphorylation of Smad2 in EDL while phosphorylated levels of Smads2 and 3 tended to be reduced in TA

Since receptors were specifically knocked out in skeletal myofibres, other cell types such as SCs, fibroblasts or inflammatory cells remained sensitive to TGF-β signalling. Therefore, the effect of receptor knockout on *Tgf-β1* and myostatin (*Mstn*) expression were determined. In TA muscles of Acvr1b^fl/fl^:Tgfbr1^fl/fl^ animals, *Tgf-β1* expression levels were 2.2-fold higher compared to those of control animals, while *Tgf-β1* expression levels of Acvr1b^fl/fl^ and Tgfbr1^fl/fl^ animals did not differ from those of control animals. *Mstn* expression levels within Acvr1b^fl/fl^:Tgfbr1^fl/fl^ mice were only increased compared to those of Acvr1b^fl/fl^ animals (Figure 3C). These results indicate that lack of both *Tgfbr1* and *Acvr1b* in skeletal myofibres resulted in increased *Tgf-β1* expression in muscle tissue, which may be associated with an enhanced local regeneration.

Expression levels of various growth factors, i.e. insulin-like growth factor 1Ea (*Igf1ea),* mechano growth factor *(Mgf),* fibroblast growth factor 2 (*Fgf-2*), hepatocyte growth factor (*Hgf*), interleukin-6 (*Il-6*) and vascular endothelial growth factor (*Vegf*) may contribute to SC proliferation or myofibre size (Arsic et al., 2004; Coleman et al., 1995; Lefaucheur & Sebille, 1995; Pedersen, Steensberg, & Schjerling, 2001; Serrano, Baeza-Raja, Perdiguero, Jardi, & Munoz-Canoves, 2008; Tatsumi, Anderson, Nevoret, Halevy, & Allen, 1998; Yang & Goldspink, 2002). No significant differences were observed in expression levels of *Igf1ea, Il-6* or *Fgf-2*. *Mgf* expression levels in TA muscle of Acvr1b^fl/fl^:Tgfbr1^fl/fl^ animals were reduced compared to those of Acvr1b^fl/fl^ or control animals. *Vegf* expression levels of Acvr1b^fl/fl^:Tgfbr1^fl/fl^ animals were reduced compared to those of all other groups, while *Vegf* levels of Tgfbr1^fl/fl^ animals were reduced compared to those of control animals. In contrast, *Hgf* expression levels of Acvr1b^fl/fl^:Tgfbr1^fl/fl^ animals were increased compared to those of all other groups (Figure 3C). These results suggest that in TA muscle of Acvr1b^fl/fl^:Tgfbr1^fl/fl^ animals enhanced expression of *Hgf* may contribute to the observed increase in SC number and myofibre hypertrophy.

Myostatin and activin A both have been shown to reduce protein synthesis by decreasing phosphorylation of Akt and its downstream target p70S6 kinase (p70S6K) (Amirouche et al., 2009; J. L. Chen et al., 2014; McFarlane et al., 2006; Trendelenburg et al., 2009). In TA muscle of Acvr1b^fl/fl^:Tgfbr1^fl/fl^ animals, phosphorylated Akt had increased by 2.3-fold compared to that of control animals. A similar but insignificant trend was observed for total Akt relative intensity. No significant differences in the phosphorylated Akt/total Akt ratio were observed. However, phosphorylated p70S6K had increased by 1.9-fold compared to that of control animals, while total p70S6K was not significantly affected. As a consequence, the phosphorylated p70S6K/total p70S6K was increased by 1.7-fold (Figure 3D). Together, these results indicate simultaneous knockout of *Acvr1b* and *Tgfbr1* in skeletal myofibre stimulates that the protein synthesis via activation of Akt/mTOR/p70S6K signalling.

Finally, myostatin and TGF-β1 have been indicated to stimulate muscle specific E3 ubiquitin ligases muscle RING-finger protein-1 (MuRF-1) and atrogin-1. *Murf-1*expression levels in TA muscles of Acvr1b^fl/fl^:Tgfbr1^fl/fl^ mice were significantly lower compared to those of Acvr1b^fl/fl^ or control animals. *Atrogin-1* expression did not differ between groups (Figure 3C). Together, these results indicate that simultaneous knockout of *Acvr1b* and *Tgfbr1* in skeletal myofibre reduces protein breakdown via suppression of *Murf-1* expression.

### Myofibre specific receptor knockout affected the inflammatory response upon acute injury

After characterisation of uninjured TA muscles, effects of receptor knockout on early TA muscle regeneration were examined two and four days after CTX acute injury. Two days post injury, the injury site was characterised by increased interstitial space, indicating degradation of the endomysium, and the presence of damaged myofibres, as can be observed as unspecific green secondary antibody staining (Bencze, Periou, Baba-Amer, & Authier, 2019). Furthermore, mononuclear cells (i.e. inflammatory cells, fibroblasts or SCs) had infiltrated the interstitial space within the injury site. Together, these observations indicate that at 2 days post injury, the inflammatory response is high and damaged myofibres have not started to regenerate yet. Four days post injury, the injury site was occupied by small, regenerating, eMyHC^+^ myofibres with centrally located nuclei. Mononuclear cells were located in the interstitial space, but the inflammatory response appears to be reduced compared to that observed at 2 days post injury (Figure 4A). No significant differences in injury size between groups were observed (Figure 4C).

**Figure 4.**
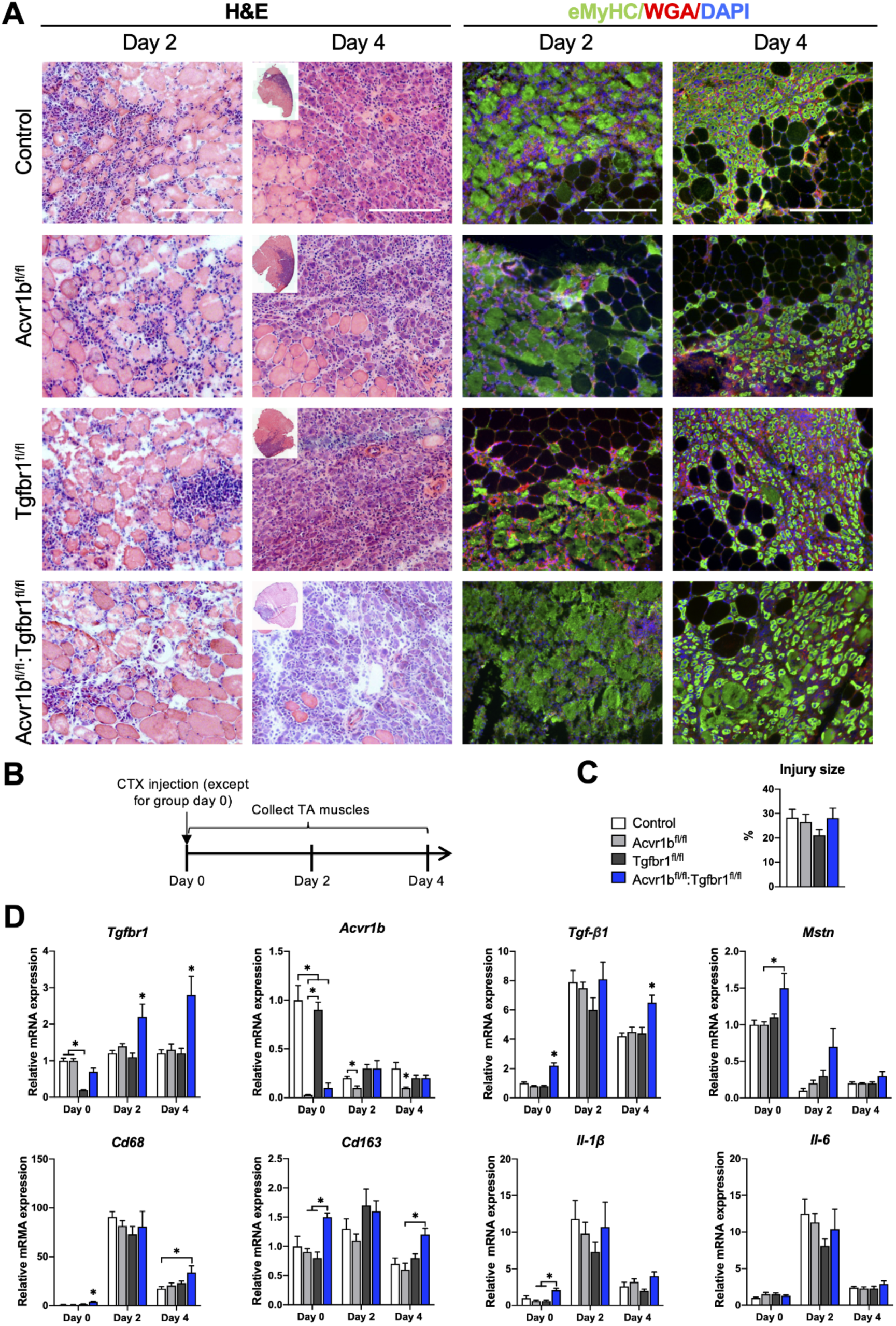
Immune response was slightly enhanced in muscle of Acvr1b^fl/fl^:Tgfbr1^fl/fl^ mice. (A) Representative images of H&E and eMyHC staining of TA sections at 2 and 4 days after CTX injection. Scale bars=250 μm. (B) Scheme shows CTX injection in TA and sample collection. (C) Percentage of injury area was not significantly different between groups. (D) Relative gene expressions in TA in the absence (day 0) or presence of CTX injection after 2 and 4 days. Results are presented as mean + SEM. N=5-8 mice, *: *P* < 0.05. Significant difference between individual groups is indicated by lines with a *. Single * indicates significant difference compared to all other groups at the same time point.

Since various cell types in muscle tissue remain sensitive to TGF-β signalling in the current model, effects of CTX injury on relative mRNA expression levels of *Tgfbr1, Acvr1b, Tgf-β1* and *Mstn* were examined. Two and four days post injury, relative *Tgfbr1* expression was increased in TA muscle of Tgfbr1^fl/fl^ and Acvr1b^fl/fl^:Tgfbr1^fl/fl^ animals compared to day 0, which suggests that *Tgfbr1* mRNA is highly expressed in mononuclear cells (i.e. fibroblasts, inflammation cells and SCs) that infiltrate the injury site. For all groups, relative *Tgf-β1* expression peaked at day 2 post injury. At day 4 post injury in Acvr1b^fl/fl^:Tgfbr1^fl/fl^ animals, *Tgf-β1* expression levels remained significantly increased compared to those of other groups (Figure 4D).

At day 2 and 4, *Acvr1b* expression in Tgfbr1^fl/fl^ and control animals decreased compared to day 0, whereas *Acvr1b* expression in Acvr1b^fl/fl^ and Acvr1b^fl/fl^:Tgfbr1^fl/fl^ animals increased. Relative *Mstn* expression levels were decreased at day 2 and 4 post injury. These data suggest that under control conditions *Acvr1b* and *Mstn* are highly expressed in skeletal myofibres, while mononuclear cells that infiltrate the injury site express relatively little *Acvr1b* and *Mstn* (Figure 4D).

TGF-β1 plays an important role in the early inflammatory response after acute muscle injury. Inflammatory cells (i.e. neutrophils and macrophages), which infiltrate damaged muscle, digest cellular debris and secrete inflammatory cytokines, such as interleukin-1β (*Il-1β*) and *Il-6*. Here, we showed that in all groups, relative mRNA levels of macrophage specific protein cluster of differentiation 68 (*Cd68*) (da Silva, Platt, de Villiers, & Gordon, 1996; M. J. Smith & Koch, 1987), *Il-1β* and *Il-6* peaked two days post injury. At day 0 and 4, in TA muscle of Acvr1b^fl/fl^:Tgfbr1^fl/fl^ animals *Cd68* expression was increased compared to all other groups or control animals, respectively. At day 0 and 4, macrophage specific *Cd163* (Schaer et al., 2001) expression levels of Acvr1b^fl/fl^:Tgfbr1^fl/fl^ animals were increased compared to those of Acvr1b^fl/fl^ or Tgfbr1^fl/fl^ animals (Figure 4D). At day 0 in Acvr1b^fl/fl^:Tgfbr1^fl/fl^ animals expression levels of *Il-1β* were increased compared to those of Acvr1b^fl/fl^ and Tgfbr1^fl/fl^ animals. Upon injury, no differences in *Il-1β* and *Il-6* expression levels were observed between groups (Figure 4D). Together, these results suggest that in this model the inflammatory response peaks approximately 2 days post injury.

### Lack of both *Acvr1b* and *Tgfbr1* stimulated CSA of regenerating myofibres, myogenic gene expression and number of differentiating muscle cells during regeneration

Effects of receptor knockout on muscle regeneration after acute injury were examined (Figure 5A). In Acvr1b^fl/fl^:Tgfbr1^fl/fl^ animals CSA of regenerating myofibres was increased compared to Acvr1b^fl/fl^ and Tgfbr1^flfl^ animals, but not compared to controls (Figure 5B). Regeneration index (RI) was reduced in muscle tissue of Acvr1b^fl/fl^ and Tgfbr1^flfl^ animals compared to that of control animals, while RI of Acvr1b^fl/fl^:Tgfbr1^fl/fl^ animals was not significantly different compared to that of other three groups (Figure 5B). Next, we hypothesized an increased immune response was involved in the accelerated muscle regeneration process after cardiotoxin induced muscle injury in the absence of Acvr1b and Tgfbr1. Macrophages were identified by F4/80 in IF staining (Figure 5-figure supplement 1). The number of macrophages in TA of Acvr1b^fl/fl^:Tgfbr1^fl/fl^ animals at day 4 post injury was significantly increased compared to that in control animals (Figure 5C). Taken together, after acute injury individual knockout of *Acvr1b* or *Tgfbr1* expression in mature myofibre reduced myofibre regeneration, while simultaneous knockout of *Acvr1b* and *Tgfbr1* stimulated this which was accompanied by increased infiltration of macrophages.

**Figure 5.**
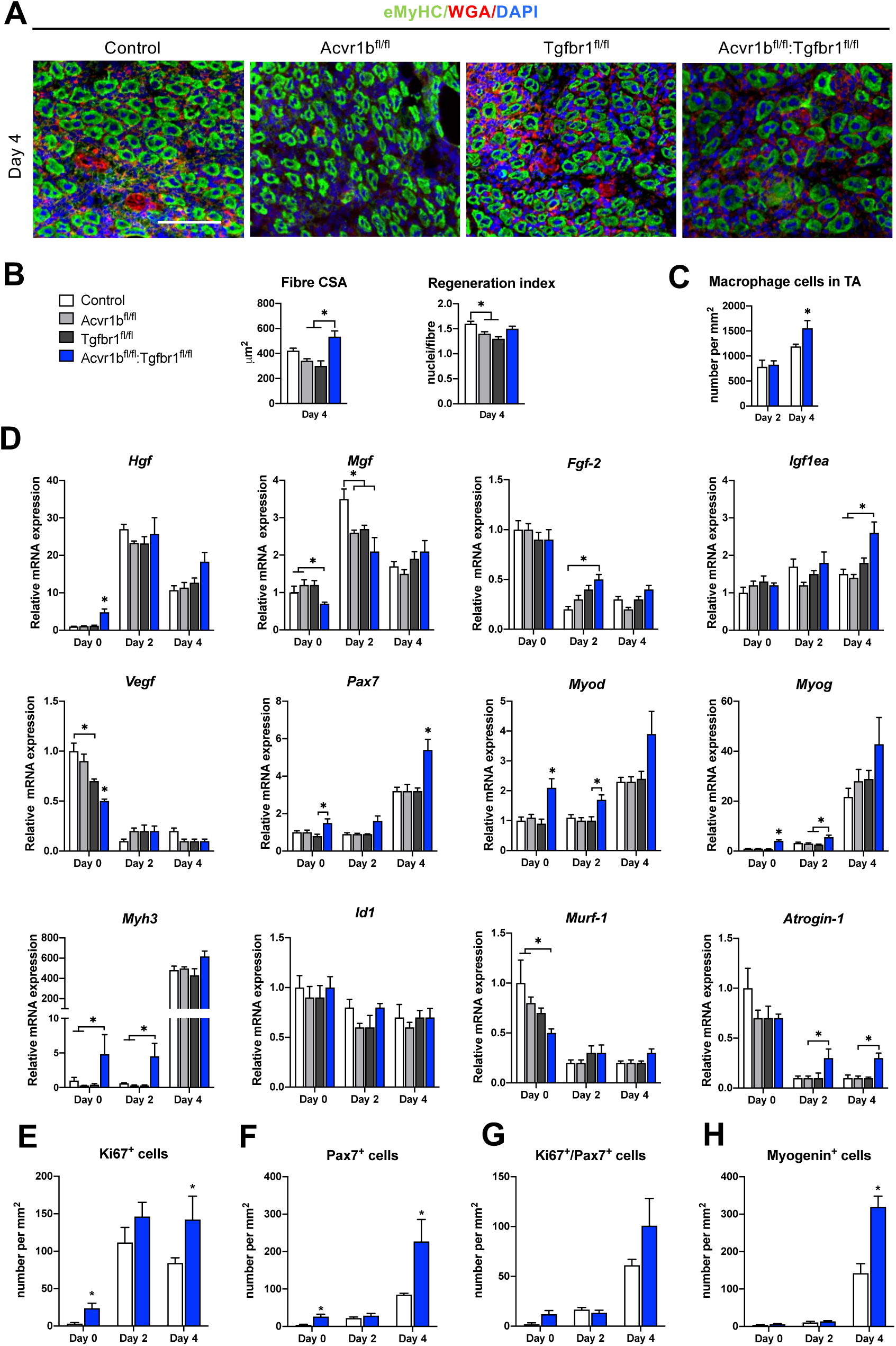
Acvr1b^fl/fl^:Tgfbr1^fl/fl^ mice showed enhanced CSA of regenerating myofibres and early enhanced expression of myogenic genes and differentiating cells after acute injury. (A) IF staining images represent eMyHC^+^ myofibres 4 days after CTX injection. Scale bar=100μm. (B) CSA of eMyHC^+^ myofibres in injured area increased in Acvr1b^fl/fl^:Tgfbr1^fl/fl^ mice compared to Acvr1b^fl/fl^ and Tgfbr1^fl/fl^ animals, while RI was decreased in both Acvr1b^fl/fl^ and Tgfbr1^fl/fl^ mice compared to controls. (C) Number of macrophages was quantified in the injured area. (D) Relative gene expression in TA in absence (day 0) or presence of CTX injection after 2 and 4 days are presented. Increased number of Ki67^+^ cells (E) and Pax7^+^ (F) cells were found in TA of Acvr1b^fl/fl^:Tgfbr1^fl/fl^ mice in absence of injury as well as 4 days after CTX injection. (G) Four days post injury, number of Ki67^+^/Pax7^+^cells was not different between control and Acvr1b^fl/fl^:Tgfbr1^fl/fl^ mice. (H) More Myogenin^+^ cells were found in injured area of Acvr1b^fl/fl^:Tgfbr1^fl/fl^ mice on day 4 post injury. Results are presented as mean + SEM. N=5-8 mice, *: *P* < 0.05. Significant difference between individual groups is indicated by lines with a *. Single * indicates significant difference compared to all other groups at the same time point.

The effects of receptor knockout on expression of genes involved in SC activation, differentiation and muscle growth were analysed in order to understand observed differences in RI and CSA of regenerating myofibres. First of all, mRNA expression levels of various growth factors were differently affected by receptor knockout. *Hgf* and *Mgf* expression peaked at day 2 post injury. At day 2 in Acvr1b^fl/fl^ and Acvr1b^fl/fl^:Tgfbr1^fl/fl^ mice *Mgf* expression was lower compared to control animals. At day 4 in Acvr1b^fl/fl^:Tgfbr1^fl/fl^ animals *Igf1ea* levels were increased compared to those of Acvr1b^fl/fl^ and control animals. *Vegf* and *Fgf-2* expression levels decreased after injury. At day 2 in Acvr1b^fl/fl^:Tgfbr1^fl/fl^ mice *Fgf-2* expression was increased (Figure 5D). Together, these results indicate that in simultaneous receptor knockout enhanced *Igf1ea* and *Fgf-2* expression post injury contribute to the accelerated early regeneration.

Proper muscle regeneration is regulated by sequential expression of myogenic genes. Thus, relative mRNA expression levels of myogenic genes (i.e., paired box protein 7 (*Pax7*), myoblast determination protein 1 (*Myod*), myogenin (*Myog*), muscle embryonic myosin heavy chain (*Myh3*) as well as inhibitor of differentiation 1 (*Id1*)), were examined. At day 4, in all groups expression of *Pax7, Myod, Myog* and *Myh3* had increased. At day 0 in Acvr1b^fl/fl^:Tgfbr1^fl/fl^ mice, *Pax7* expression was increased compared to that of Tgfbr1^fl/fl^ mice, while at day 4 in Acvr1b^fl/fl^:Tgfbr1^fl/fl^ mice *Pax7* expression was increased compared to that in all other groups. At day 0 in Acvr1b^fl/fl^:Tgfbr1^fl/fl^ mice *Myod* and *Myog* expression levels were increased compared to those in other groups, while at day 2 in Acvr1b^fl/fl^:Tgfbr1^fl/fl^ mice *Myod* expression levels were increased compared to those in Tgfbr1^fl/fl^ mice and *Myog* expression levels were increased compared to those in both Acvr1b^fl/fl^ and Tgfbr1^fl/fl^ animals. At day 4 post injury, no differences in *Myod* or *Myog* expression levels were observed between groups, although a trend suggested that expression of both genes was increased in Acvr1b^fl/fl^:Tgfbr1^fl/fl^ mice. At both day 0 and day 2, in Acvr1b^fl/fl^:Tgfbr1^fl/fl^ mice *Myh3* levels were increased compared to those in both Acvr1b^fl/fl^ and control animals. At day 4 post injury, no differences in *Myh3* expression were observed between groups (Figure 5D). Receptor knockout did not affect *Id1* expression. Taken together, these results show that in TA myofibre specific *Acvr1b* and *Tgfbr1* receptor knockout stimulates myogenic gene expression.

Lastly, relative expression levels of muscle specific E3 ligases MuRF-1 and atrogin-1 during early regeneration were considered. MuRF-1 and atrogin-1 are expressed in mature myofibres, but not in SCs and inhibit myofibre growth and hypertrophy. For Acvr1b^fl/fl^, Tgfbr1^fl/fl^ and control animals, at day 4 *Murf-1* levels were decreased compared to those at day 0, while for Acvr1b^fl/fl^:Tgfbr1^fl/fl^ animals no significant differences in *Murf-1* expression were observed over time. At both day 2 and 4 post injury, in Acvr1b^fl/fl^:Tgfbr1^fl/fl^ animals *Atrogin-1* expression levels were relatively increased compared to those in Acvr1b^fl/fl^ mice, while no differences in *Murf-1* expression levels were observed between groups (Figure 5D). Together, these results indicate E3 ligases do not play a role in the observed increase in size of regenerating myofibres.

On day 0, the number of proliferating cells (Ki67^+^) in low oxidative region of TA of Acvr1b^fl/fl^:Tgfbr1^fl/fl^ animals was about 7.6-fold higher than that in control animals (Figure 5E). Two days after injury, a 6-fold increase of proliferating cells was found in Acvr1b^fl/fl^:Tgfbr1^fl/fl^ animals compared to that on day 0. Moreover, at day 4 after injury, the number of proliferating cells in Acvr1b^fl/fl^:Tgfbr1^fl/fl^ animals was 1.7-fold higher than that in control animals. To determine whether the increased CSA of regenerating myofibres in Acvr1b^fl/fl^:Tgfbr1^fl/fl^ animals was due to an increased SCs number and advanced differentiation of myoblasts, we tested SCs proliferation and activation status. Although at day 0 and 4, the number of SCs (Pax7^+^) cells was more in Acvr1b^fl/fl^:Tgfbr1^fl/fl^ animals compared to that in control animals (Figure 5F), the number of proliferating SCs (Ki67^+^/Pax7^+^) did not differ from that in control (Figure 5G, Figure 5-figure supplement 2,). Nevertheless, an accelerated rate of increase in Ki67^+^/Pax7^+^ cells was shown. Note that, at day 4 after injury the number of myogenin^+^ cells was more than 2.2-fold higher in Acvr1b^fl/fl^:Tgfbr1^fl/fl^ animals (Figure 5H, Figure 5-figure supplement 3). These findings indicate that muscle regeneration upon acute injury was improved in Acvr1b^fl/fl^:Tgfbr1^fl/fl^ animals, which was attributed to an accelerated myogenic process.

### Simultaneous knockout of both *Acvr1b* and *Tgfbr1* within the myofibre enhanced ECM deposition

Another essential aspect of muscle regeneration is connective tissue remodelling. Figure 5A shows Sirius Red stainings at different stages of regeneration. At day 0, myofibres were surrounded by a thin layer of endomysium. At 2 days post injury, this endomysium appears to be disrupted for a large part. At day 4 post injury, a large amount of connective tissue may be observed surrounding the regenerating myofibres (Figure 6A).

**Figure 6.**
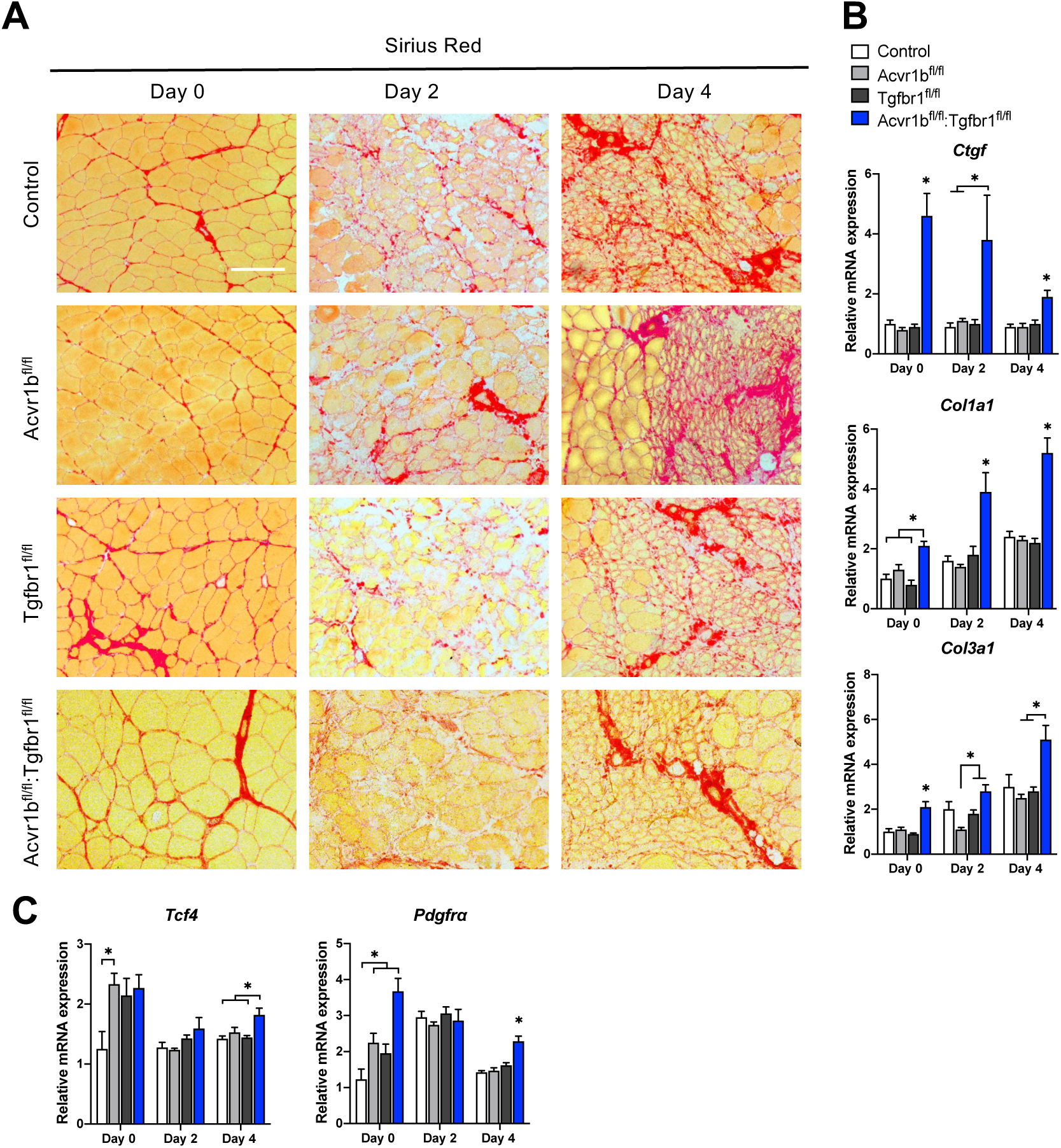
Relative mRNA expression levels of ECM components were enhanced in Acvr1b^fl/fl^:Tgfbr1^fl/fl^ mice. (A) Sirius Red staining shows collagen deposition in absence (day 0) or presence of CTX injection after 2, and 4 days (scale bar=100μm). (B, C) Relative gene expression in TA muscle in absence (day 0) or presence of CTX injection after 2 and 4 days. Results are presented as mean + SEM. N=5-8 mice, *: *P* < 0.05. Significant difference between individual groups is indicated by lines with a *. Single * indicates significant difference compared to all other groups at the same time point.

Effects of myofibre specific *Acvr1b* and *Tgfbr1* receptor knockout on ECM remodelling were assessed by examining connective tissue growth factor (*Ctgf*), collagen type 1, alpha 1 (*Col1a1*) and collagen type 3, alpha 1 (*Col3a1*) expression. At all times, in Acvr1b^fl/fl^:Tgfbr1^fl/fl^ animals *Ctgf* and *Col1a1* mRNA expression levels were substantially increased compared to those of control animals or all groups. At day 0, in Acvr1b^fl/fl^:Tgfbr1^fl/fl^ animals, *Col3a1* expression levels were increased compared to those in other groups. At day 2 and 4, in Acvr1b^fl/fl^:Tgfbr1^fl/fl^ animals *Col3a1* expression was increased compared to Acvr1b^fl/fl^ or both Acvr1b^fl/fl^ and Tgfbr1^fl/fl^ animals (Figure 6B).

To determine whether the number of fibroblasts was increased in TA muscles by knockout of *Acvr1b* and *Tgfbr1*, we assessed relative mRNA expression levels of fibroblast markers, transcription factor 4 (*Tcf4*) and platelet-derived growth factor receptor A (*Pdgfra*) (Mathew et al., 2011), were determined. At day 0, *Tcf4* expression levels of TA in Acvr1b^fl/fl^ mice were increased compared to those in control animals, but were not different from those in Tgfbr1^fl/fl^ or Acvr1b^fl/fl^:Tgfbr1^fl/fl^ animals. In Acvr1b^fl/fl^:Tgfbr1^fl/fl^ animals, relative *Pdgfra* expression levels were increased compared to those of control and Acvr1b^fl/fl^ animals. Noteworthy, four days post injury, both *Tcf4* and *Pdgfra* mRNA levels were increased in Acvr1b^fl/fl^:Tgfbr1^fl/fl^ animal compared to those in control mice (Figure 6C).

## Discussion

The aim of this study was to investigate effects of mature myofibre specific knockout of type I receptors *Tgfbr1* and *Acvr1b* on muscle morphology as well as early muscle regeneration, inflammation and collagen deposition in both uninjured muscle tissue and after acute CTX injury. We observed that simultaneous knockout of *Acvr1b* and *Tgfbr1* resulted in a substantial increase in TA and EDL muscle mass as well as type IIB myofibre CSA. *Tgfbr1* knockout only marginally increased muscle mass, while *Acvr1b* knockout did not affect muscle mass. In the low oxidative region of TA muscle tissue of Acvr1b^fl/fl^:Tgfbr1^fl/fl^ animals the percentage of type IIB myofibres was reduced, while in EDL no differences in myofibre type distribution were observed. Remarkably, simultaneous knockout of both *Acvr1b* and *Tgfbr1* caused spontaneous regeneration and an increase in SC number in the low oxidative region of TA and in EDL muscle tissue. Lack of both *Acvr1b* and *Tgfbr1* in the skeletal myofibre of TA increased myofibre CSA of regenerating myofibres, number of regenerating cells and macrophages during regeneration, as well as enhanced expression levels of myogenic gene expression and growth factors. In both uninjured and regenerating muscles, simultaneous knockout of *Acvr1b* and *Tgfbr1* in the myofibre resulted in increased ECM and fibroblasts gene expression. Figure 7 shows a schematic summarising the main results.

**Figure 7.**
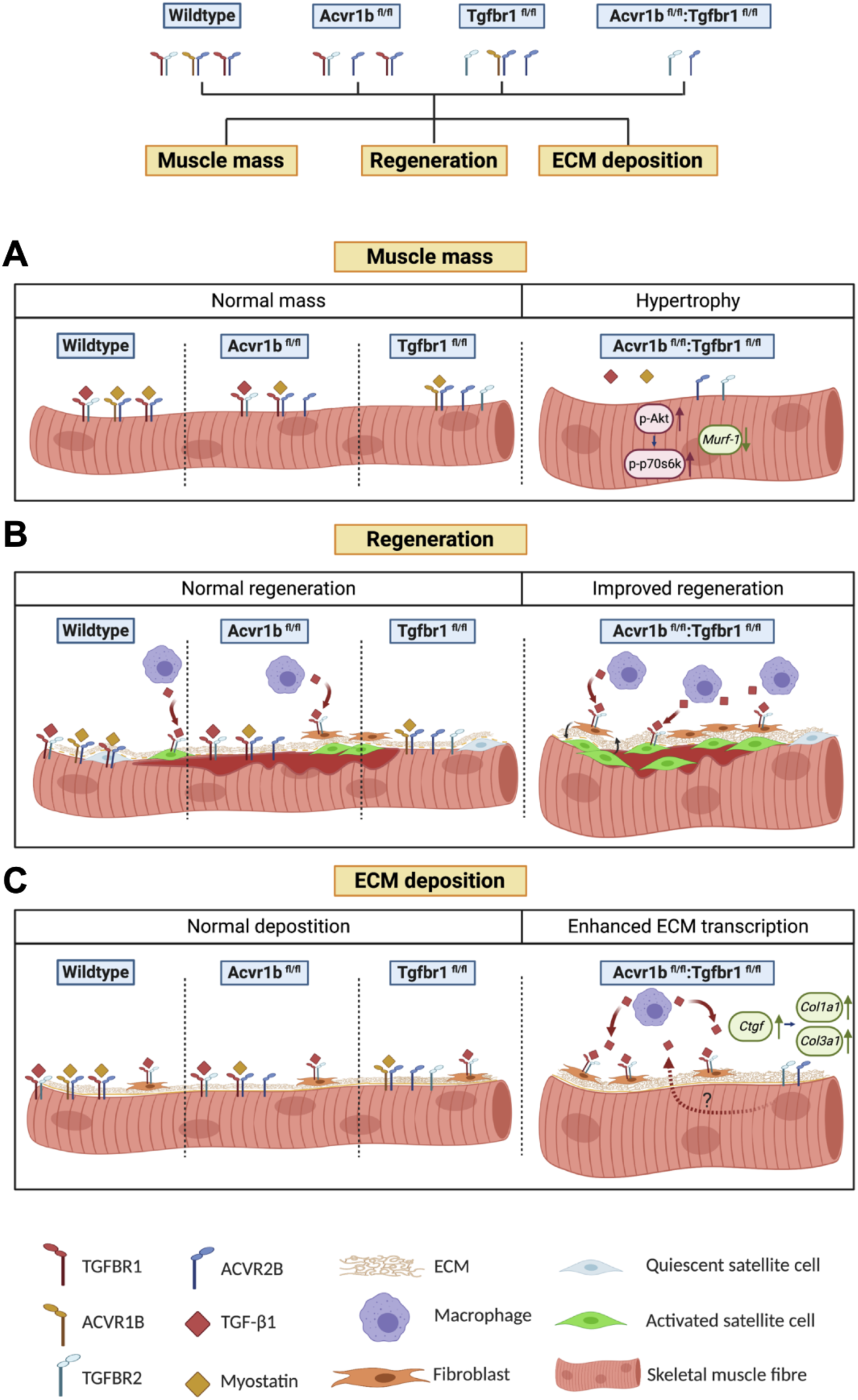
Schematic diagram of the effects of single or combined muscle specific knockout of *Tgfbr1* and/or *Acvr1b* receptors on muscle hypertrophy, regeneration and expression of ECM components. (A) Myofibre size is not affected after individual knockout of *Acvr1b* or *Tgfbr1*, which indicates that these receptors have redundant effects on muscle size and that myostatin signals via both receptors to control muscle mass. Simultaneous knockout of both *Acvr1b* and *Tgfbr1* inhibits signalling of TGF-β, myostatin and activin A and stimulates protein synthesis via the Akt/mTOR/p70S6K pathway, while inhibiting protein breakdown through repression of *Murf-1* levels, resulting in substantial muscle hypertrophy. (B) Upon acute injury, simultaneous knockout of combined *Acvr1b* and *Tgfbr1* accelerates early muscle regeneration, as observed by increased myogenic gene expression as well as increased CSA of regenerating myofibres. An increased number of SCs likely contributes to these effects. (C) Simultaneous myofibre specific knockout of *Acvr1b* and *Tgfbr1* induces mRNA expression of ECM components. These effects are likely caused by enhanced TGF-β1 signalling in fibroblasts. Schematic is created using BioRender.

Together, these results indicate that simultaneous receptor knockout stimulates muscle hypertrophy and promotes early muscle regeneration upon injury, whereas individual receptor knockout does not.

### Simultaneous knockout of *Acvr1b* and *Tgfbr1* induces hypertrophy by both inhibiting protein degradation and stimulating protein synthesis

Our data show that simultaneous myofibre specific knockdown of both *Acvr1b* and *Tgfbr1* is required for muscle hypertrophy, while inhibition of *Acvr1b* does not affect muscle mass or myofibre size and inhibition of *Tgfbr1* has only marginal effects. Supporting our data, a recent study showed that simultaneous inhibition of *Acvr1b* and *Tgfbr1* was required to enhance muscle mass, while individual receptor inhibition had little effect (Lee et al., 2020). These results indicate that *Acvr1b* and *Tgfbr1* have redundant functions in the regulation of muscle mass.

Myostatin and activin A negatively regulate muscle mass by stimulating protein degradation through upregulation of E3 ligases and reducing protein synthesis through a decrease in phosphorylation of Akt and its downstream target p70S6K (Amirouche et al., 2009; J. L. Chen et al., 2014; McFarlane et al., 2006; Trendelenburg et al., 2009). TGF-β overexpression *in vivo* has been suggested to increase atrogin-1 expression and concomitantly cause muscle atrophy (Mendias et al., 2012). Simultaneous receptor knockout increased relative expression of phosphorylated Akt and lack of both *Acvr1b* and *Tgfbr1* increased the phosphorylated p70S6K/total p70S6K ratio, which indicates increased protein synthesis via the Akt/mTOR/p70S6K pathway contributes to muscle hypertrophy.

Simultaneous knockout of *Acvr1b* and *Tgfbr1* also reduced mRNA expression levels of E3 ligases *Murf-1*, indicating reduced protein degradation. Simultaneous knockout of *Acvr1b* and *Tgfbr1* increased *Hgf* expression levels. HGF signalling has been shown to protect skeletal muscle against atrophy after denervation or in muscle pathology by reducing relative *Murf-1* and *Atrogin-1* expression levels (Choi, Lee, Lee, Ko, & Kim, 2018). Together, these results indicate that simultaneous knockout of both *Acvr1b* and *Tgfbr1* both decrease protein breakdown and stimulate protein synthesis.

The TGF-β type I receptor knockout induced an increase in Akt signalling and reduction in E3 ligase expression are likely mediated at least in part via elevated *Hgf* expression levels in myofibres. In both myoblasts and myotubes, HGF stimulates the Akt/mTOR pathway (A. L. Chen et al., 2012; Perdomo et al., 2008). In addition, reduced Smad2/3 phosphorylation within myofibres likely contributes to the increase in Akt signalling and reduction in E3 ligase expression (Goodman, McNally, Hoffmann, & Hornberger, 2013; Sartori et al., 2009). In TA and EDL of Acvr1b^fl/fl^:Tgfbr1^fl/fl^ animals, Smad2/3 phosphorylation was or tended to be reduced, respectively. Note that TGF-β type I receptors were specifically knocked out in myofibres and that Smad2/3 in various other cell types can still be phosphorylated by TGF-β1, myostatin and activin A, masking the changes in myofibres.

TGF-β1, myostatin and activin A regulate skeletal muscle mass via similar mechanisms. Previous research has shown that inhibition of myostatin and to a lesser extent activin A is sufficient to induce muscle hypertrophy (Wu et al., 2017). Here we show that inhibition of TGF-β1 or activin A signalling via their type I receptor is insufficient to induce muscle hypertrophy. In muscle myostatin likely signals via both type I receptors to regulate muscle mass. In muscles that lack either *Acvr1b* or *Tgfbr1,* we observed no changes in Smad2/3 signalling, which is in accordance with the observation there is no or modest effect on muscle hypertrophy. Targeting both receptors is indispensable to substantially reduce TGF-β1/myostatin/activin A signalling and induce muscle hypertrophy.

### Simultaneous knockout of *Acvr1b* and *Tgfbr1* specifically enhances type IIB myofibre CSA without accretion of myoblasts

Simultaneous knockout of *Acvr1b* and *Tgfbr1* most substantially increased type IIB myofibre CSA. This is likely the result of myofibre type-related hypertrophic capacity rather than myofibre type-specific receptor knockout bias (McCarthy et al., 2012b). It has been suggested that mainly fast twitch myofibres possess the ability to hypertrophy, while slow twitch myofibres are unlikely to increase in size (van Wessel et al., 2010). Additionally, ACVR2B is more abundantly expressed in type II than type I myofibres, thus a more substantial effect on myofibre hypertrophy was expected upon type I receptor depletion (Babcock, Knoblauch, & Clarke, 2015). Remarkably, type IIB myofibre hypertrophy occurred without apparent accretion of myonuclei, which resulted in an approximately 70% increase in myonuclear domain. The lack of difference in the number of myonuclei per myofibre cross-section together with the lack of difference in myonuclear length in type IIB myofibres of TA indicates that the total number of nuclei per myofibre was not affected by simultaneous knockout of *Acvr1b* and *Tgfbr1*. Although e.g. exercise induced hypertrophy is often accompanied by increased myonuclei number (Conceicao et al., 2018; van der Meer, Jaspers, Jones, & Degens, 2011), the myonuclear domain is known to be flexible and increases in myonuclear domain of 30% have been reported (Murach, Englund, Dupont-Versteegden, McCarthy, & Peterson, 2018). Moreover, inhibition of myostatin signalling using soluble ACVR2B leads to hypertrophy without accretion of SCs (Lee et al., 2012). Here, we show that in type IIB myofibres myonuclear domain can increase by at least 70%, without requirement of accretion of myonuclei to sustain myofibre growth. To the best of our knowledge such increase in myonuclear domain has not been reported before.

Since myonuclei are required for mitochondrial biogenesis, a local reduction in oxidative capacity was expected (Hock & Kralli, 2009; Kotiadis, Duchen, & Osellame, 2014). In this study, knockout of both *Acvr1b* and *Tgfbr1* resulted in decreased SDH activity in the low oxidative region of TA. Previous research has shown that an inverse relation exists between myofibre CSA and oxidative capacity, whereas myofibre CSA is positively correlated to glycolytic capacity (Rivero, Talmadge, & Edgerton, 1998). Recent evidence suggests that, similar to “Warburg effect” in tumours, in hypertrophying skeletal muscle reprogramming towards a more glycolytic metabolism occurs. Glycolytic enzyme pyruvate kinase muscle isoform 2 (PKM2), which is particularly highly expressed in type II myofibres, contributes to the increased hypertrophic potential of type II myofibres (Verbrugge et al., 2020).

Although lack of *Acvr1b* and *Tgfbr1* decreased SDH activity, integrated SDH activity (SDH activity times CSA) was increased. Previous research has shown that integrated SDH activity correlates with the maximal rate of oxygen consumption (VO_2max_) and mitochondrial density, which suggests that the total oxidative capacity of these myofibres in Acvr1b^fl/fl^:Tgfbr1^fl/fl^ animals was increased (W. J. van der Laarse, Diegenbach, & Elzinga, 1989). Present data show that by targeting both receptors simultaneously it is possible to deviate from the tight relation between myofibre size and oxidative metabolism (i.e. simultaneous increases in both myofibre size and oxidative capacity). The role of both receptors in the synthesis of mitochondria warrants further investigation.

### Lack of both *Acvr1b* and *Tgfbr1* reduces the percentage of type IIB myofibres within the low oxidative region of the TA

Another remarkable finding was the reduction in the percentage of type IIB myofibres in the low oxidative region of the TA muscle of Acvr1b^fl/fl^:Tgfbr1^fl/fl^ animals. In the high oxidative region of the TA muscle as well as the EDL, no differences in myofibre type distribution were observed. In contrast to our findings, previous research has shown increased percentage of fast, glycolytic myofibres in skeletal muscle of Mstn^-/-^ mice (Amthor et al., 2007; Girgenrath, Song, & Whittemore, 2005; Hennebry et al., 2009). Moreover, increased myostatin/activin A signalling in follistatin mutant mice showed increased the percentage of slow, oxidative myofibres (Lee et al., 2010). Note that in a genetic knockout mouse model, absence of myostatin precedes myogenesis and may influence skeletal muscle development, whereas in our model TGF-β signalling was inhibited in mature skeletal muscle. The reduction in type IIB myofibres may also be a consequence of local damage to the myofibres, rather than a phenotypical change caused by receptor knockout. Taken together, these results indicate type I receptor knockout in mature myofibres has minor effects on myofibre type distribution.

### Simultaneous knockout of *Acvr1b* and *Tgfbr1* in myofibre may result in accelerated early regeneration

We observed that simultaneous knockout of *Acvr1b* or *Tgfbr1* increased the CSA of regenerating myofibres compared to individual receptor knockout, while a trend was visible compared to controls. In addition, at day 0, simultaneous receptor knockout enhanced *Hgf* expression as well as the number of SCs per myofibre and concomitantly relative expression levels of *Pax7* and *Myod,* indicating that SCs were activated prior to CTX injection. Muscle regeneration is dependent on activation of Pax7^+^ SCs and sequential expression of myogenic genes (Charge & Rudnicki, 2004; Delaney, Kasprzycka, Ciemerych, & Zimowska, 2017; Ishido & Kasuga, 2011; Lepper, Partridge, & Fan, 2011). HGF is the primary growth factor for SC activation and may have accelerated early muscle regeneration (Allen, Sheehan, Taylor, Kendall, & Rice, 1995; Gal-Levi, Leshem, Aoki, Nakamura, & Halevy, 1998; Miller, Thaloor, Matteson, & Pavlath, 2000; Tatsumi et al., 1998). Receptor knockout did not affect *Hgf* expression after injury, which indicates HGF expression and subsequent SC activation is likely not induced by lack of TGF-β signalling in the myofibre, but rather a consequence of spontaneous damage and regeneration.

Moreover, 2 or 4 days post injury simultaneous knockout of *Acvr1b* and *Tgfbr1* increased expression levels of *Fgf-2* or *Igf1ea*. Simultaneous overexpression of IGF-1 and FGF-2 has a synergistic effect on both myoblast proliferation as well as fusion index (Allen & Boxhorn, 1989). Thus, simultaneous knockout of *Acvr1b* and *Tgfbr1* in skeletal myofibre likely enhances early muscle regeneration via enhanced expression of *Igf1ea* and *Fgf-2.* However, myogenic gene expression was increased prior to *Igf1ea* and *Fgf-2* upregulation, which indicates that although these growth factors may positively contribute to early regeneration, upregulation of *Igf1ea* and *Fgf-2* cannot fully explain effects on early regeneration.

Simultaneous receptor knockout did not reduce mRNA expression of E3 ligases during regeneration, indicating that the increase in myofibre CSA is not caused by reduced protein breakdown. In contrast, during regeneration *Atrogin-1* expression is enhanced in Acvr1b^fl/fl^:Tgfbr1^fl/fl^ animals compared to other groups, but here the enhanced *Atrogin-1* levels may correspond with muscle regeneration.

Moreover, for Acvr1b^fl/fl^:Tgfbr1^fl/fl^ animals the numbers of SCs and differentiating myoblasts within the injured regions 4 days post injury were more than doubled compared to those in control animals. The observation that at all time points the number of proliferating SCs (i.e. Ki67^+^/Pax7^+^ cells) was not different between control and Acvr1b^fl/fl^:Tgfbr1^fl/fl^ animals indicates that in the Acvr1b^fl/fl^:Tgfbr1^fl/fl^ animals muscle damage had initiated activation and proliferation in the injured region between day 2 and 4. Alternatively, SCs had migrated from adjacent myofibres to the site of injury or from intact regions along the myofibres (Ishido & Kasuga, 2011; Schultz, Jaryszak, & Valliere, 1985). The accelerated myoblast proliferation and differentiation likely contributed to the enhanced protein synthesis and hypertrophy of newly formed myofibres.

A limitation of this study is that we did not observe later stages of muscle regeneration. Additional research is required to determine whether changes observed in this study ultimately result in a shorter regeneration period.

### *Acvr1b* and *Tgfbr1* affect inflammatory response after injury

Proper activation of the immune response and expression of inflammatory cytokines is important for myoblast proliferation and myogenic gene expression during early muscle regeneration (Cantini et al., 1995; Chaweewannakorn et al., 2018; Grabiec et al., 2013; Zhang et al., 2013). In absence of injury, compared with control, 8912310-fold more number of macrophages was found the in low oxidative area of TA in Acvr1b^fl/fl^:Tgfbr1^fl/fl^ animals, indicating increased immune cells residence at baseline before CTX injury. Two days post injury a large infiltration of mononucleated cells was observed in all groups, as well as a peak in relative expression levels of *Tgf-β1*, *Cd68*, *Il-1β* and *Il-6*.

Macrophages play an important role in regulation of muscle regeneration (Tidball, 2017). Macrophages are classified in M1 (pro-inflammatory) and M2 (anti-inflammatory) macrophages (Mosser & Edwards, 2008). Early after injury, gene expression of pan-macrophages marker *Cd68* and M2 macrophage marker *Cd163* (Hu et al., 2017), as well as the number of macrophages were increased in both control and Acvr1b^fl/fl^:Tgfbr1^fl/fl^ animals. Expression levels *Il-6* and *Il-1β*, which are typical cytokines expressed by M1 macrophages, were not higher than in control animals, while *Igf-1ea* expression, also known to be expressed by M1 macrophages, was increased in Acvr1b^fl/fl^:Tgfbr1^fl/fl^ animals which was likely advantageous to expand the SC pool and to induce hypertrophy of newly formed myofibres. Moreover, TGF-β1 expression was increased in muscle with simultaneous knockout of Acvr1b^fl/fl^:Tgfbr1^fl/fl^. At a later stage after injury, M2 macrophages are known to promote myogenic differentiation and stimulate ECM deposition by releasing TGF-β1 (Arnold et al., 2007; Novak, Weinheimer-Haus, & Koh, 2014). Taken together, we conclude that muscle-specific lack of both receptors promotes an inflammatory response by enhanced infiltration of macrophages which is associated with accelerated muscle regeneration.

### Lack of both *Acvr1b* and *Tgfbr1* enhances gene expression of ECM components in both intact and injured TA muscle

In both uninjured TA muscle tissue, as well as after CTX injury simultaneous knockout of *Tgfbr1* and *Acvr1b* in skeletal myofibre increased *Ctgf*, *Col1a1* and *Col3a1* mRNA expression. These increases in gene expression were conceivably caused by TGF-β signalling in other cell types present within the muscle tissue, i.e. fibroblasts. This hypothesis is supported by the infiltration of fibroblasts in TA muscle of Acvr1b^fl/fl^:Tgfbr1^fl/fl^ animals in the absence of injury, as well as increased gene expression levels of *Pdgfrα*. Furthermore, after injury in Acvr1b^fl/fl^:Tgfbr1^fl/fl^ animals *Tgfbr1, Tcf4* and *Pdgfrα* levels were increased compared to those in groups, which indicated increased infiltration of non-muscle cells upon injury. Together these results support the hypothesis that TGF-β and myostatin act on fibroblasts and possibly other cell types within the muscle tissue to induce expression of collagens. We previously showed that inhibition of *Tgfbr1* in C2C12 myoblasts reduced *Ctgf* and *Col1a1* expression (Hillege et al., 2020). Moreover, systemic administration of anti-TGF-β or soluble ACVR2B in murine X-linked muscular dystrophy (mdx) mice reduced muscular fibrosis (Andreetta et al., 2006; Bo Li, Zhang, & Wagner, 2012). In conclusion, to reduce expression of ECM components within skeletal muscle inhibition of TGF-β signalling in other cell types such as fibroblasts and satellite cells is required.

Chronic excessive ECM deposition leads to increased muscle stiffness and loss of function. However, transiently enhanced ECM deposition is essential to early muscle regeneration and results in scar free muscle repair in various types of acute injury (Hardy et al., 2016; Mahdy, Lei, Wakamatsu, Hosaka, & Nishimura, 2015). In this study after CTX injury *Col1a1* and *Col3a1* expression increased in all groups. Transient enhanced ECM deposition is required to maintain muscle structural integrity and provides a scaffold for regenerating myofibres (Kaariainen, Jarvinen, Jarvinen, Rantanen, & Kalimo, 2000). Furthermore, interaction between fibroblasts and SCs appears to be essential for proper muscle regeneration, since fibroblasts prevent early differentiation of SCs, while in turn SCs control the number of fibroblasts (Murphy, Lawson, Mathew, Hutcheson, & Kardon, 2011). Thus the observed enhanced expression of ECM components in TA that lacks both receptors may contribute to the increased number of SCs at day 0 and acceleration of early muscle regeneration.

### Implications for ACVR1B and TGFBR1 inhibition as potential therapeutic strategy

An important limitation of our study is that we are investigating effects of *Acvr1b* and *Tgfbr1* knockout on early regeneration after an acute injury. In contrast to our model, a dystrophic or aged mouse model has characteristics such as chronic inflammation, impaired regeneration and fibrosis. Further research is required to determine how *Acvr1b* and *Tgfbr1* knockout affects long term regeneration capacity, chronic inflammation and fibrosis in a pathological model.

Our data indicate that simultaneous knockout of *Acvr1b* and *Tgfbr1* in the mature myofibre causes muscle hypertrophy and accelerates early muscle regeneration. The inflammatory response is likely independent on TGF-β signalling within the myofibre. Furthermore, specifically targeting *Tgfbr1* and *Acvr1b* in mature myofibre actually increased relative ECM gene expression levels. This effect is likely the result of enhanced TGF-β signalling in other cell types (i.e. fibroblasts and inflammatory cells). Thus, targeting TGF-β signalling in immune cells and fibroblasts present in muscle tissue is likely required to alleviate chronic inflammation and fibrosis.

Taken together, our data indicate that individually inhibiting either *Acvr1b* or *Tgfbr1* may not be sufficient to alleviate muscle pathologies, nevertheless combined inhibition of both *Acvr1b* and *Tgfbr1* may increase muscle mass and accelerate early muscle regeneration.

## Materials and Methods

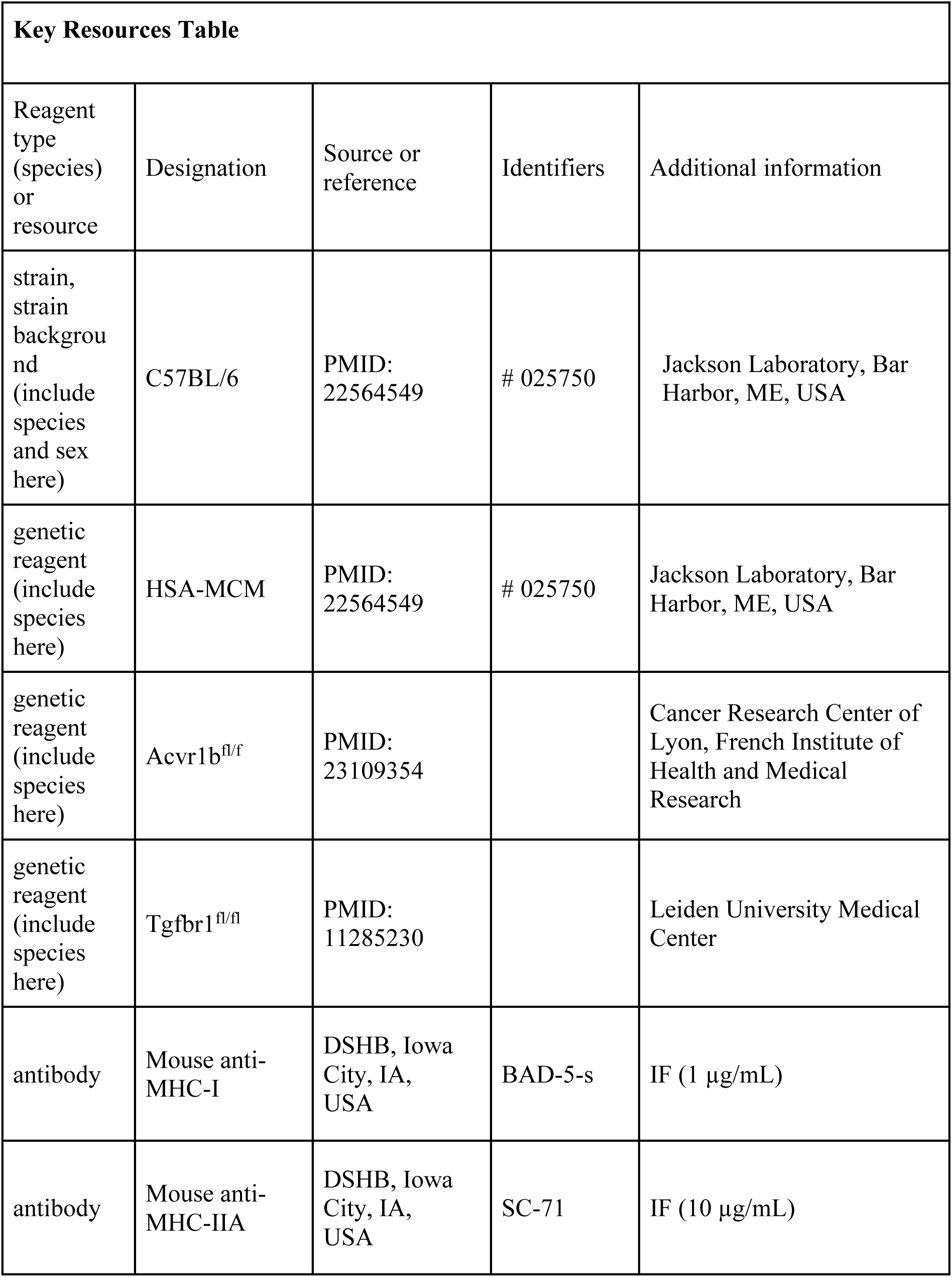
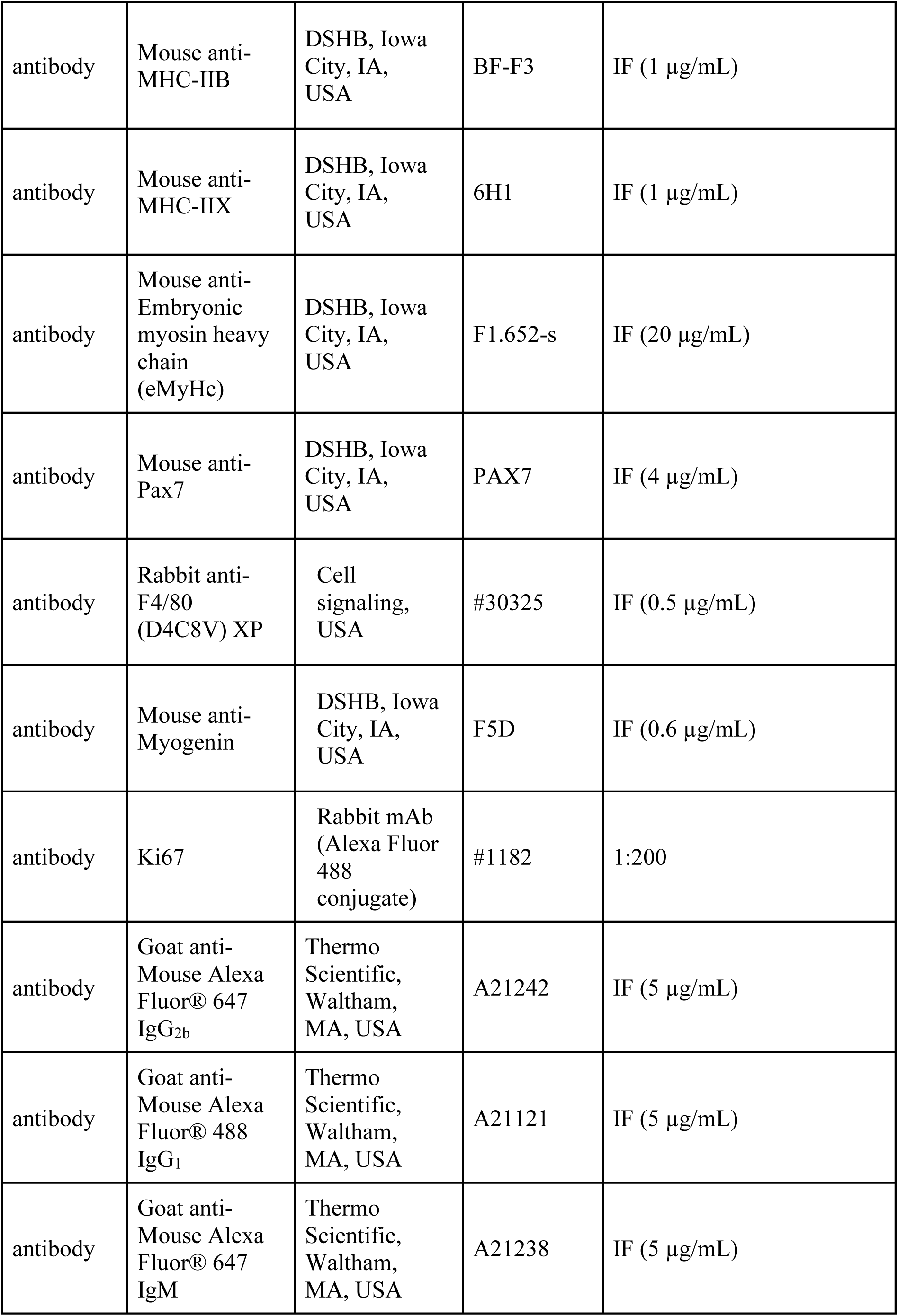
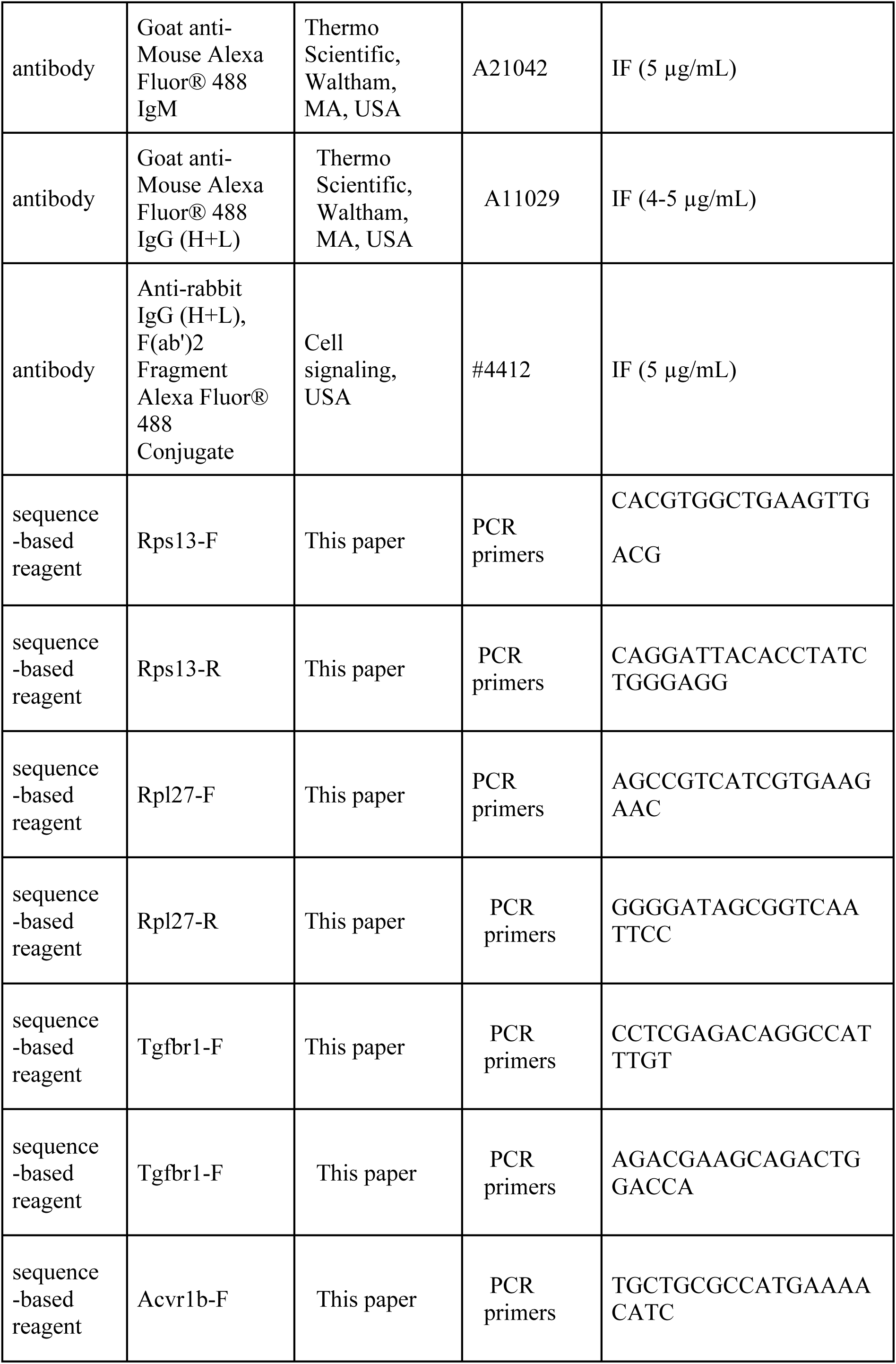
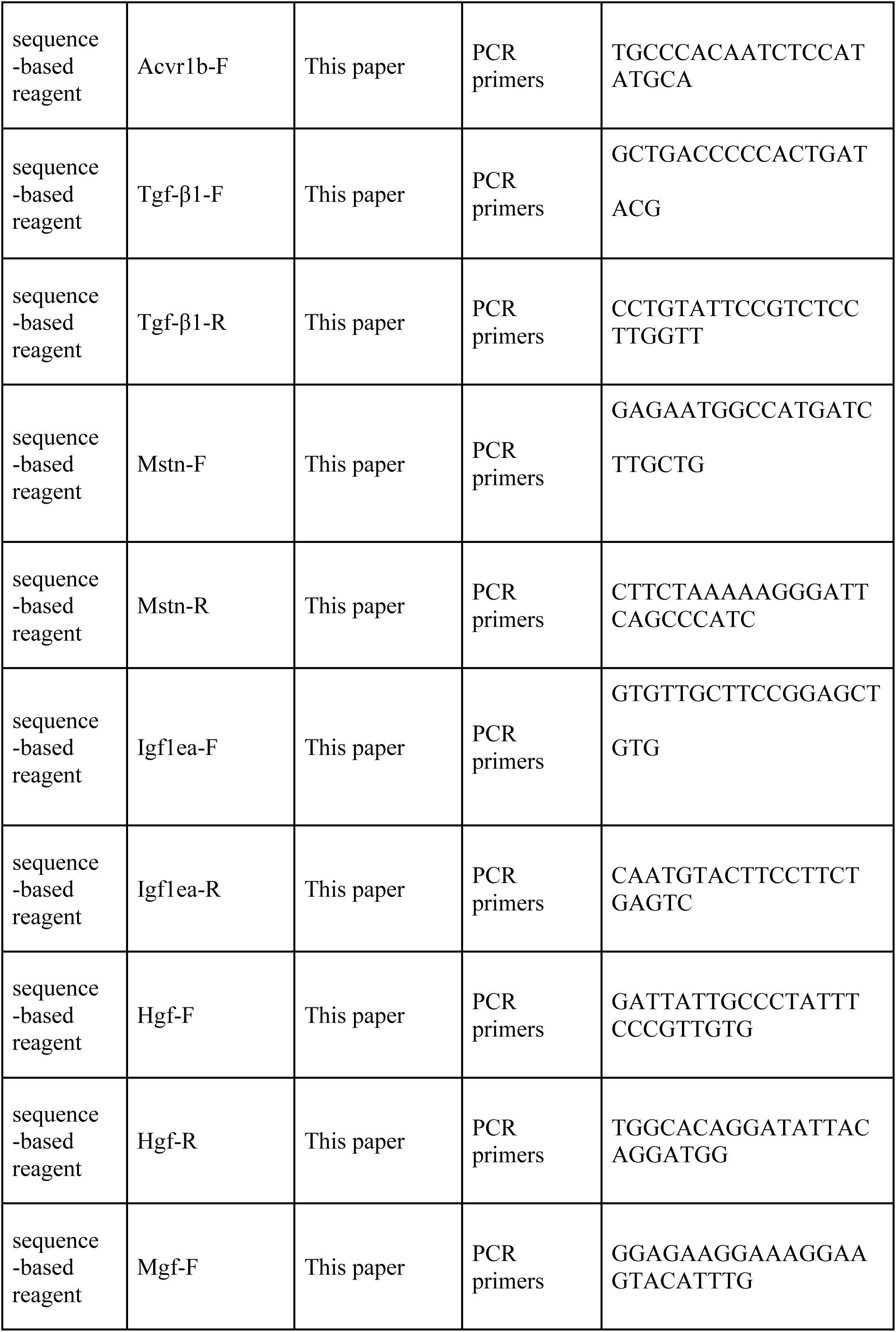
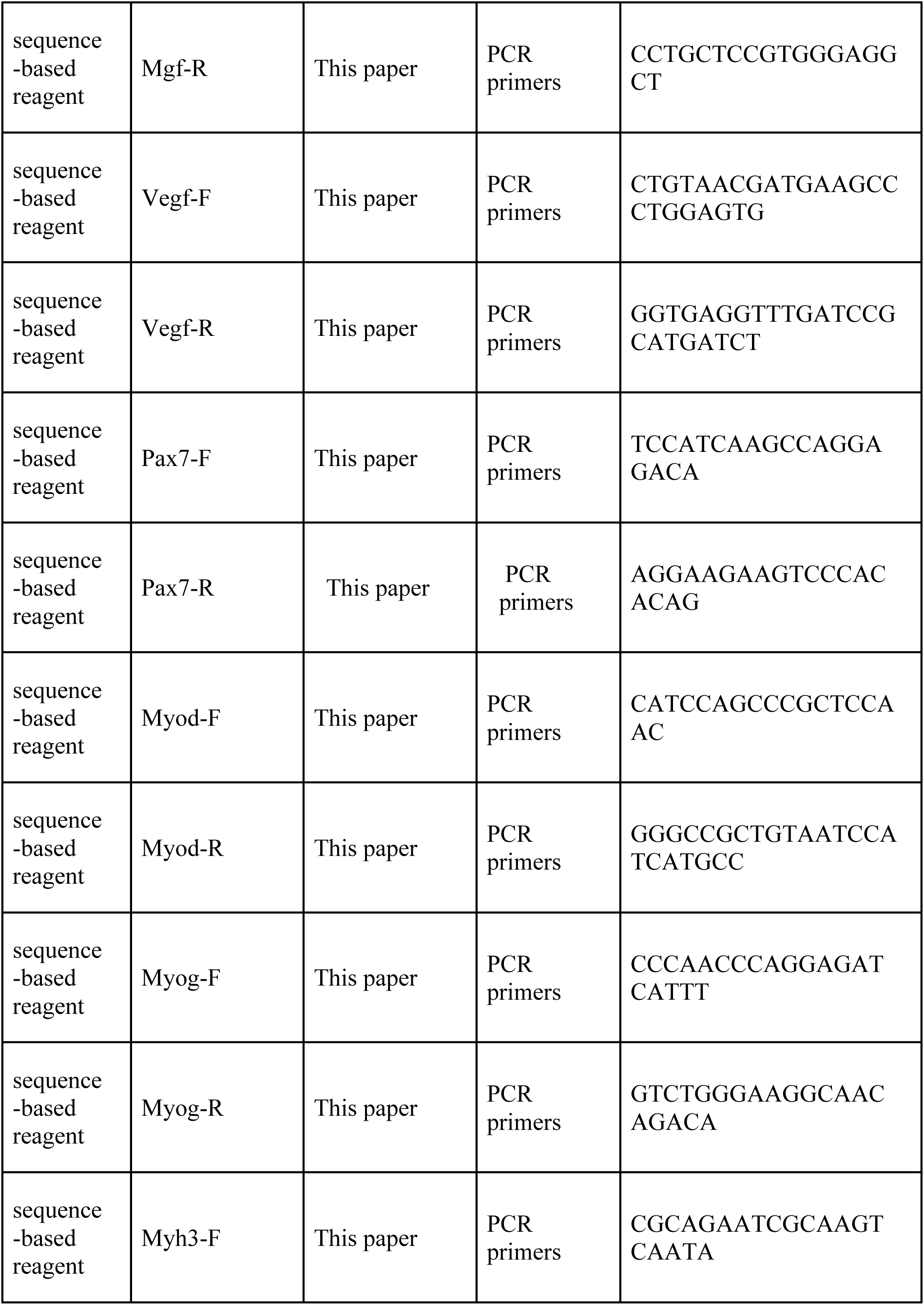
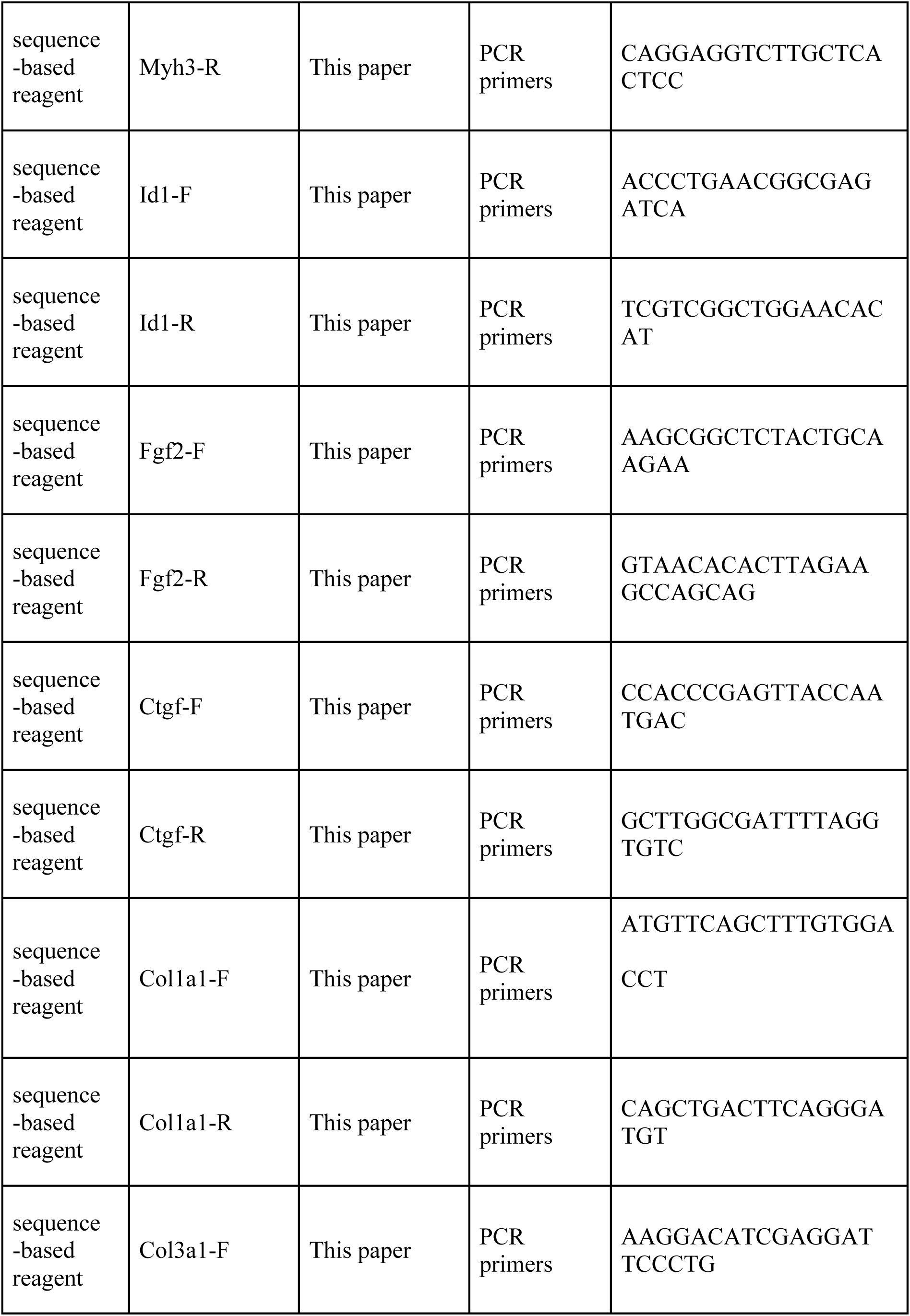
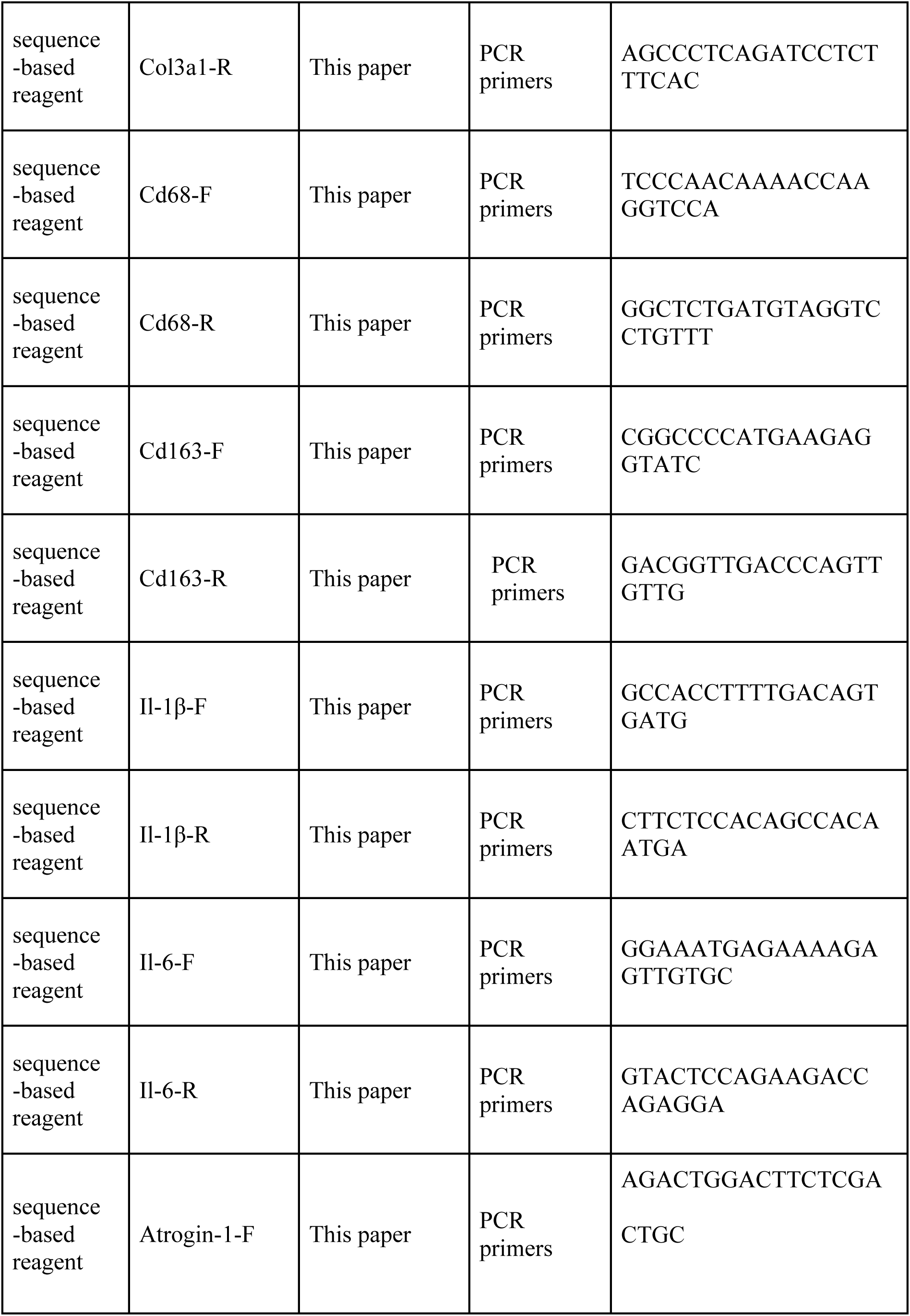
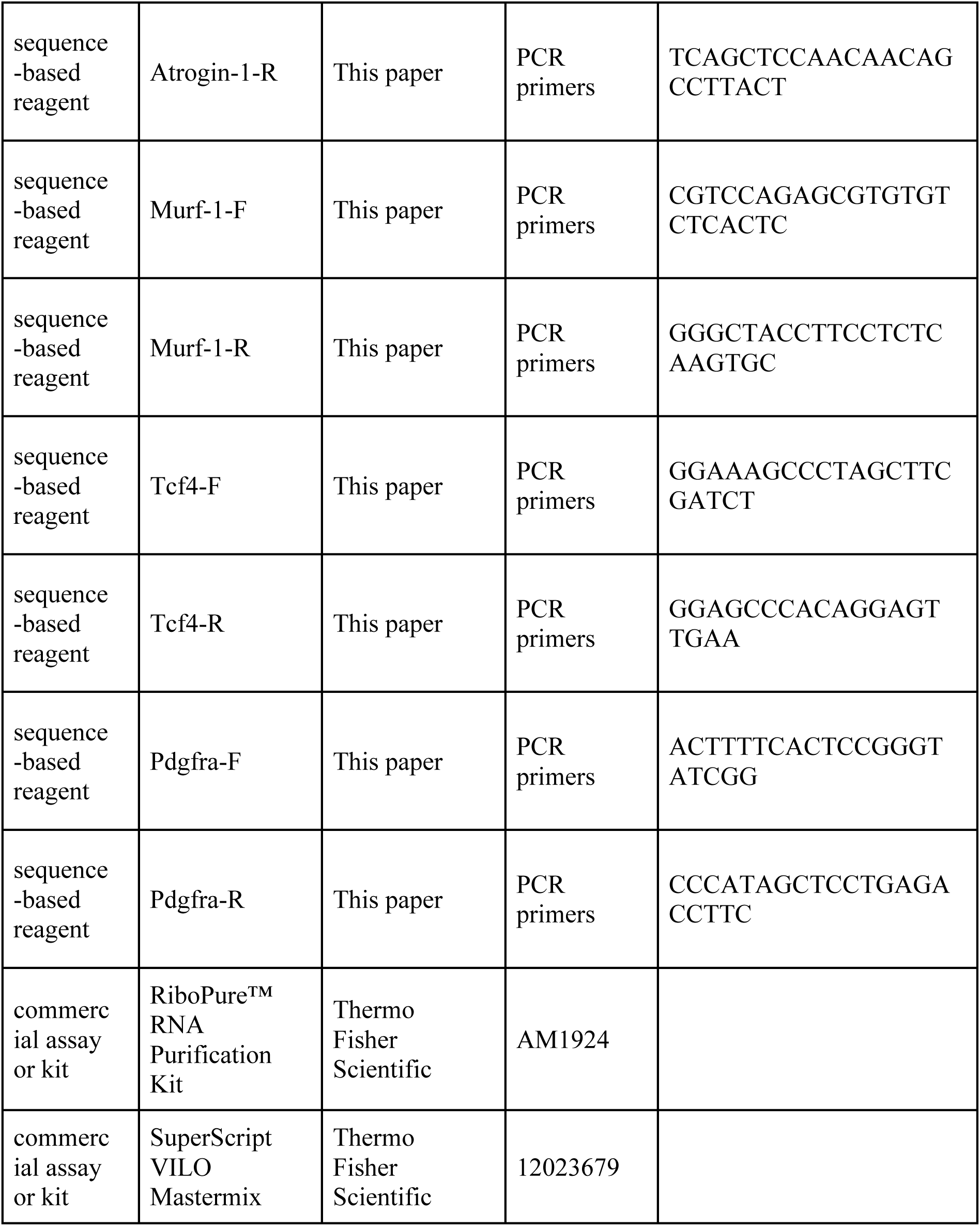

### Animal housing and welfare

The HSA-MCM transgenic mouse line (McCarthy et al., 2012b) was obtained from Jackson Laboratory, Bar Harbor, ME,USA (stock number # 025750), the Acvr1b^fl/fl^ mouse line (Ripoche et al., 2013) was obtained from Philippe Bertolino (Cancer Research Center of Lyon, French Institute of Health and Medical Research) and the Tgfbr1^fl/fl^ mouse line (Larsson et al., 2001) was provided by Peter ten Dijke (Leiden University Medical Center). All mouse lines were of a C57BL/6 background. Mouse lines were cross-bred in house to obtain mouse lines HSA-MCM:Acvr1b^fl/fl^, HSA-MCM:Tgfbr1^fl/fl^ and HSA-MCM:Acvr1b^fl/fl^:Tgfbr1^fl/fl^. Animals were housed in a controlled 12 hours light-dark cycle (light on 6:00-18:00 GMT + 1h) with a temperature of 21 ± 1°C and a humidity between 40 and 70%. Food (Teklad, Envigo, Horst, The Netherlands) and water were available at libitum. All experiments were performed according to the national guidelines approved by the Central Committee for Animal Experiments (CCD) (AVD112002017862) and the Institute of Animal Welfare (IvD) of the Vrije Universiteit Amsterdam.

### Genotyping

In these mouse models, skeletal myofibre specific Cre expression is driven by the HSA promoter which can be activated by TMX, resulting in the deletion of targeted exon of 5 and 6 of *Acvr1b* and exon 3 of *Tgfbr1* resulting in a targeted knockout of the gene. Genotyping for the HSA-MCM, Acvr1b^fl/fl^ and Tgfbr1^fl/fl^ genes was performed by isolating DNA from ear biopsies of offspring and PCR was performed in a 2720 thermal cycler (Applied Biosystems, Foster City, CA, USA). PCR master mix per sample was prepared by mixing 0.2 μl of AmpliTaq Gold DNA polymerase, 2.5 μl of gold buffer, 1.5 μl of MgCl_2_ (Thermo Fisher Scientific, 4311806, Waltham, MA, USA), 0.5 μl of dNTPs (100mM diluted 10×, Invitrogen 10297018, Carlsbad, CA, USA), 1 μl of each primer diluted in DNAse/RNAse free water to obtain a volume of 23 μl Master mix per sample. 2 μl DNA was added per sample. The following PCR programs were used: for HSA-MCM and Acvr1b^fl/fl^: 94°C for 5 minutes, followed by a 35× cycle of 94°C for 30 seconds, 58°C for 30 seconds and 72°C for 10 minutes, finishing with 72°C for 10 minutes and cooled down to 4°C. PCR program for Tgfbr1^fl/fl^: 94°C for 4 minutes, followed by a 35× cycle of 94°C for 30 seconds, 50°C for 45 seconds and 72°C for 1 minute, finishing with 72°C for 5 minutes and cooled down to 4°C. Amplified DNA was mixed with 5 μl loading buffer and samples were loaded in a 4% agarose gel using SYBR safe DNA gel staining 1000× concentrate (Thermo Fisher Scientific s33102, Waltham, MA, USA), DNA was separated by electrophoresis (25 minutes, 75 V) and gel image was taken using an Image Quant LAS 500 chemo luminescence CCD camera (GE healthcare, life sciences, Chicago, IL, USA).

Primer sequences: HSA-MCM gene: Forward, 5′- GCATGGTGGAGATCTTTGA-3′ (McCarthy et al., 2012b) and Reverse, 5′-CGACCGGCAAACGGACAGAAGC-’3 (McCarthy, Srikuea, Kirby, Peterson, & Esser, 2012a). Acvr1b^fl/fl^ gene: Acvr1b In4, 5’- CAGTGGTTAAGAACACTGGC-3’, Acvr1b In5, 5’- GTAGTGTTATGTGTTATTGCC -3’ And Acvr1b In6, 5’GAGCAAGAGTTTCTCTATGTAG-3’ (Ripoche et al., 2013). Tgfbr1^fl/fl^ gene: Forward, 5’- CCTGCAGTAAACTTGGAATAAGAAG-’3, reverse, 5’- GACCATCAGCTGTCAGTACCC-3’ (Protocol 19216: Standard PCR Assay - Tgfbr1<tm1.1Karl>, Jackson Laboratory, Bar Harbor, ME,USA).

### Cardiotoxin induced injury assay

Animals of each genotype were assigned to each timepoint randomly to ensure that groups were on average the same age at the time of the first TMX injection. Littermates were assigned to different timepoints. Six weeks old HSA-MCM:Acvr1b^fl/fl^, HSA-MCM:Tgfbr1^fl/fl^, HSA-MCM:Acvr1b^fl/fl^:Tgfbr1^fl/fl^ and HSA-MCM Cre^+^ male mice were injected intraperitoneally with 100 mg/kg/day tamoxifen (Sigma-Aldrich, T5648, Saint-Louis, MO, USA) in sunflower oil (10 mg/ml) for five consecutive days. Five weeks post TMX injections, mice were injected intramuscularly in the TA muscles of both hind limbs with 20 µl CTX from Naja pallida (Latoxan Laboratory, L8102, Portes les Valence, France) in phosphate buffered saline (PBS) (10 µM). CTX was slowly injected (1 µl/s) into mid muscle belly using a Hamilton syringe with attached 34G needle inserted in a 15-25° angle, 2-3 mm deep. Mice were shortly anesthetised using isoflurane 1.5-3% on a warm blanket during the injections. Mice were divided into three groups: mice that were sacrificed 2 days or 4 days post injury and mice with no CTX injection, that were sacrificed at day 0, which functioned as a baseline control. Mice were sacrificed by cervical dislocation and TA muscles were isolated and frozen in isopentane cooled in liquid nitrogen. Each subgroup contained 5-8 mice.

### RNA isolation and reverse transcription

Whole TA muscle was used for RNA isolation. 50 mg cryopreserved TA muscle was homogenised (Potter S 8533024, B. BRAUN) in 700 µl TRI reagent (Invitrogen, 11312940, Carlsbad, CA, USA) and incubated at room temperature (RT) for 5 minutes. Samples were centrifuged for 10 minutes (4°C, 12000 g). Supernatant was transferred to a new tube and 70 µl bromochloropropane (Sigma-Aldrich, B9673, Saint Louis, MO, USA) was added. Lysates were inverted and incubated at RT for 5 minutes and centrifuged (4°C, 12000 g, 10 minutes). RNA containing supernatant was transferred to a new centrifuge tube and washed with 100% ethanol 2:1. RNA was further isolated using the RiboPure RNA purification kit (Thermo Fisher Scientific, AM1924, Waltham, MA, USA). Then, 500 ng RNA and 4 µl SuperScript VILO Mastermix (Invitrogen, 12023679, Carlsbad, CA, USA) were diluted to 20 µl in RNAse free water and reverse transcription was performed in a 2720 thermal cycler (Applied Biosystems, Foster City, CA, USA), using the following program: 10 minutes at 25 °C, 60 minutes at 42 °C and 5 minutes at 85 °C. cDNA was diluted 10× in RNAse free water.

### Quantitative real time PCR

5 μl of PowerUp SYBR Green master mix (Applied Biosystems, A25742, Foster City, CA, USA), 3 μl of primer mix and 2 μl of cDNA were added in duplo in a 96 wells plate. The program ran on the Quant Studio 3 real time PCR (Applied Biosystems, Foster City, CA, USA) was 2 minutes at 50°C, 2 minutes at 95°C, 40× 1 second at 95°C and 30 seconds at 60°C, 15 seconds at 95°C, 1 minute at 60°C and 15 seconds at 95°C. Geometric mean of reference genes ribosomal protein S13 (*Rps13*) and ribosomal protein L27 (*Rpl27*) was used to correct for cDNA input. The efficiency of all primers sequences (Key Resources Table) was > 98%.

### Western blot

Fifty 20 μm cross sections of TA and EDL muscles were obtained using a cryostat microtome (Microm HM550, Adamas Instruments, Rhenen, The Netherlands). GM tissue and whole EDL muscles were lysed (Potter S 8533024, B. BRAUN) in RIPA buffer (Sigma-Aldrich, R0278, Saint Louis, MO, USA) containing 1 tablet of protease inhibitor (Sigma-Aldrich, 11836153001) and 1 tablet of phosStop (Sigma-Aldrich, 04906837001) per 10ml. The total protein concentration in the lysates was determined using a Pierce BCA Protein Assay kit (Thermo Scientific, 23225). A 4–20% Mini-PROTEAN® TGX™ Precast Protein Gels was used. 15µl sample mix containing 12µg total protein and 4µl sample buffer (4× Laemmli Sample Buffer, Bio-Rad, 1610747) with 10% mercaptoethanol (Bio-Rad, 1610710) was heated to 95⁰C for 5 minutes, cooled on ice and loaded onto the gel. After electrophoresis, proteins were transferred onto a polyvinylidene fluoride (PVDF) membrane (GE Healthcare, 15269894) for blotting at 80V for 60 minutes. The membrane was incubated for 1 hour at RT in 2% enhanced chemiluminiscence (ECL) prime blocking agent (GE Healthcare, RPN418). Membranes were incubated overnight at 4⁰C in blocking buffer (4% bovine serum albumin (BSA) in tris-buffered saline with 0.1% Tween 20 detergent (TBST) ) with primary antibody at a dilution of 1:500 for Phospho-Smad2 (Ser465/467) (138D4) (Rabbit mAb, 3108, Cell signaling, USA), 1:500 Phospho-Smad3 (Ser423/425) (C25A9) (Rabbit mAb, 9520, Cell signalling, USA), 1:500 for Purified mouse anti-Smad2/3 (610843, BD Biosciences, USA), 1:1000 for Phospho-AKT (Ser473) (Rabbit mAb, 4060, Cell signaling, USA), 1:2000 for AKT (pan) (C67E7) (Rabbit mAb, 4691, Cell signaling, USA), Phospho-p70S6 Kinase (Thr389) (108D2) (Rabbit mAb, 8209, Cell signaling, USA), p70S6 Kinase (49D7) (Rabbit mAb, 2708, Cell signaling, USA) and Pan-Actin (Rabbit Ab, 4968, Cell signalig, USA). Incubation with secondary antibody, anti-Rabbit IgG-POD, (P0448, Agilent Dako, USA) or Rabbit anti-Mouse IgG, IgM (H+L), HRP conjugate (31457, ThermoFisher Scientific, USA) at a dilution of 1:2000 was done for 1 hour at RT in blocking buffer and detection was done using ECL detection kit (RPN2235, GE Healthcare, USA). Images were taken by the ImageQuant LAS500 (GE healthcare, life sciences, USA) and relative intensity of protein bands was quantified using ImageJ (Schneider, Rasband, & Eliceiri, 2012). Pan-Actin was used as a loading control.

### Tissue cross sectioning for histological analysis

For histological analysis, 10 μm thick cross-sections of TA or EDL muscles were obtained using a cryostat microtome (Microm HM550, Adamas Instruments, Rhenen, The Netherlands), mounted on microscope slides (super frost plus, Thermo Scientific, J1800AMNZ, Landsmeer, The Netherlands) and stored at -80°C for further analyses. In addition, for TA muscle, 10 μm thick longitudinal sections were obtained to measure the myonuclear length.

### Histochemistry staining of H&E and Sirius Red staining

For H&E staining, slides with muscle sections were air dried for 10 minutes. Dried slides were then stained in Hematoxylin for 2 minutes, then washed under tap water for 15 minutes and further stained in Eosin for 1 minute. Immediately after Eosin staining, slides were rinsed 2× in 100% alcohol for 10 seconds and 2× in xylene for 10 minutes. Stained sections were mounted in Entellan (Merck, 107960, Darmstadt, Germany), covered with coverslip and dried overnight before analysis.

For Sirius Red staining, sections were air dried for 10 minutes at RT, then they were fixed in acetone at -20°C for 10 minutes and subsequently fixed in saturated Bouin for 30 minutes at RT. Samples were stained by Sirius Red for 30 minutes and washed in 0.01 M hydrochloric acid. Then, sections were rapidly dehydrated in 100% ethanol and xylene twice. Stained sections were mounted in Entellan, covered with coverslip and dried overnight before analysis (de Bruin, Smeulders, Kreulen, Huijing, & Jaspers, 2014).

Images were captured on a Zeiss Axioskop microscope (Carl Zeiss BV, Breda The Netherlands) with Basler camera (Basler AG, Ahrensburg, Germany) using Manual WSI scanner software (Microvisioneer) for collecting whole scan images. Regeneration area percentage at day 0 was referred to area of myofibres with central nuclei divided by total area of a whole muscle cross-section. Morphometry was performed using ImageJ software (Schneider et al., 2012).

### Immunofluorescence staining

Microscope slides with muscle sections were air dried for 10 minutes. For Pax7 staining, sections were fixed with 4% paraformaldehyde (PFA) at RT for 10 minutes and washed in PBS-T (0.05% Tween 20). Sections were blocked at RT with 10% normal goat serum (NGS) (Thermo Fisher Scientific, 50062Z, Waltham, MA, USA) (60 minutes for <u>myosin heavy chain</u> (MyHC), eMyHC, F4/80 and Myogenin staining, or 30 minutes for Pax7 staining) or 5% NGS with 0.3% Triton-X100 for 60 minutes for Ki67/Pax7 staining. Sections for Pax7 staining were subsequently blocked in 1% BSA in PBS for 30 minutes. Sections were incubated in primary antibodies (Key Resources Table) in 10% NGS (for MyHC, eMyHC or Myogenin) or in 0.1% BSA (Pax7, F4/80 or Ki67/Pax7) at RT for 60 minutes or overnight. Then sections were incubated in secondary antibodies (Key Resources Table) in 10% NGS for MyHC and eMyHC staining, or in 0.1% BSA for Pax7 or F4/80 staining at RT in the dark for 60 minutes. After this, sections were incubated in 1:50 diluted wheat germ agglutinin (WGA) (Fisher Scientific, 11590816, Pittsburgh, PA, USA) in PBS at RT for 20 minutes. Finally, slides were carefully dried and mounted with Vectashield hardset mounting medium with 4‘,6-diamidino-2-phenylindole (DAPI) (Brunschwig, H1500, Amsterdam, The Netherlands). Slides were dried overnight at 4°C.

### Immunofluorescence microscopy and analysis

Images of all immunofluorescence assays were captured using a fluorescent microscope (Zeiss Axiovert 200M, Hyland Scientific, Stanwood, WA, USA) with a PCO SensiCam camera (PCO, Kelheim, Germany) using the program Slidebook 5.0 (Intelligent Imaging Innovations, Göttingen, Germany). The images were analysed using ImageJ (Schneider et al., 2012). Individual images were taken across the entire cross-section and assembled into a composite panoramic image. For TA, the inner part of the muscle tissue was referred to as high oxidative region and the outer part was referred to as low oxidative region. For myofibre type analysis, 250 myofibres within each part were characterised in TA. Hybrid myofibre fluorescence was assessed by myofibres stained by double colours and pure myofibres were determined by single colour staining (Bloemberg & Quadrilatero, 2012). To determine CSA per myofibre type, CSA was measured of 30 myofibres per type, or as many as were present within the tissue per TA. For EDL, myofibre type and CSA of all myofibres within the muscle were determined by SMASH (L. R. Smith & Barton, 2014). Within images stained for Pax7, the number of Pax7^+^ nuclei and the number of myofibres were determined in the low oxidative region of TA from about 200 myofibres (5 fields per specimen). SCs were defined as Pax7^+^ cells that were located between plasma and basal lamina of myofibre (Lindstrom & Thornell, 2009). Within muscle sections images stained for F4/80, the density of macrophages in TA at day 0 was determined in 10 randomly selected locations in TA at day 0 and at least 3 locations within the injured region at day 2 and 4 after injury. Macrophages were defined as F4/80^+^ cells that were located in the interstitial region of the myofibres at day 0. To measure the number of myonuclei per type IIB myofibre, 100 type IIB myofibres in the low oxidative region were taken into account and nuclei were considered as myonuclei when they were located within the cytoplasm below the basal lamina. To determine myonuclear length, images were taken at 10 × magnification using a fluorescent microscope. Mean myonuclear length was determined as the average value of 30 nuclei. For regeneration analysis, myofibre CSA measurement was performed by outlining 50 eMyHC^+^ myofibres from three randomly chosen fields within the regenerating area in muscle cross-sections of day 4. RI was defined as the number of nuclei within eMyHC^+^ myofibres divided by the number of eMyHC^+^ myofibres, all eMyHC^+^ myofibres in three randomly chosen fields (1.38mm^2^ per field) within the regenerating area were included in the analysis. The densities of Ki67^+^, Pax7^+^, Ki67^+^/Pax7^+^ and Myogenin^+^ cells in TA were determined by counting the number of cells per mm^2^ of muscle CSA. Ten images in low oxidative region of TA on day 0, 2 and 4 were randomly selected. All analyses were performed at 20× magnification.

### SDH assay

Succinate dehydrogenase activity was quantified according to Van der Laarse (1989). Breiefly, freshly muscle cross-sections (10 µm thick) were air dried for 15 minutes at RT. Sections were incubated in prewarmed SDH (37.5 mM NaH_2_PO_4_·H_2_O, 37.5 mM Na_2_HPO_4_·2H_2_O, added acid to pH 7.6, 75 mM sodium succinate, 5 mM NaN_3_, 0.5 g/L tetranitroblue tetrazolium (TNBT)) medium for 10 minutes at 37°C. Sections were rinsed 3 seconds in 0.01 M hydrochloric acid. Then sections were rinsed for 1 minute twice in ultrapure water. Finally, sections were mounted in glycerine gelatin (Merck, 48723, Darmstadt, Germany) with coverslips and dried overnight before analysis. Images were captured by a Leica DMRB microscope (Wetzlar, Germany) with calibrated grey filters and a CCD camera (Sony XC77CE, Towada, Japan) connected to a LG-3 frame grabber (Scion, Frederick, MD, United States). The absorbances of the SDH-reaction product in the sections were determined at 660 nm using a calibrated microdensitometer and ImageJ (Schneider et al., 2012). SDH activity (ΔA_660_ · µm^-1^ ·s^-1^) was measured by the rate of absorbance per section thickness per second ( Δ A_660_/(10 µm · 600s)) after subtracting background activity in 5 locations in low oxidative region at 10× magnification. The integrated SDH activity (ΔA_660_ · µm ·s^-1^) was defined as SDH activity × myofibre CSA. Absorbance was measured in a total of 50 myofibres per TA.

### Statistical analysis

Graphs were made in Prism version 8 (GraphPad software, San Diego, CA, USA). All data were presented as mean + standard error of the mean (SEM). Statistical analysis was performed in SPSS version 26 (IBM, Amsterdam, The Netherlands). Sample size was determined a priori, based on previously published *in vivo* research on TGF-β and myostatin. Statistical significance for multiple comparisons was determined by two-way analysis of variance (ANOVA), three-way ANOVA or independent t-test. Significance was set at *P* < 0.05. Data normality was tested with a Shapiro-Wilk test (*P* < 0.05). Homogeneity of variance was tested with a Levene’s test (*P* < 0.01). If necessary, data were square or log transformed. Post hoc Bonferroni or Games-Howell corrections were performed. If normality was violated a Kruskall Wallis test was performed.

### Data availability

This study includes data deposited in Dryad.

## Acknowledgements

This research was funded by the Prinses Beatrix Spierfonds, grant number W.OR14-17 and a grant from the China Scholarship Council (CSC grant number 201808440351). We thank Philippe Bertolino (Cancer Research Center of Lyon, French Institute of Health and Medical Research) and Peter ten Dijke (Leiden University Medical Center) for providing the Acvr1b^fl/fl^ and Tgfbr1^fl/fl^ mouse lines. We thank animal caretakers of the Universitair Proefdier Centrum of the Vrije Universiteit Amsterdam, our colleague Wendy Noort and students Elke Schmitz and Bijee Visbeek for their contribution to data analysis.

## Author contributions

Conceptualization and Methodology: M.M.G.H., A.S., R.A.G., W.M.H.H., R.T.J.

Formal Analysis: M.M.G.H., A.S.

Funding acquisition: R.T.J, W.M.H.H.

Investigation: M.M.G., A.S., R.A.G.

Resources: R.T.J., P.B.

Supervision: G.W., R.T.J.

Visualization and Writing – original draft: M.M.G.H., A.S.

Writing – review & editing: M.M.G.H., A.S., R.A.G., W.M.H.H., R.T.J.

---

**Figure 5-figure supplement 1.**
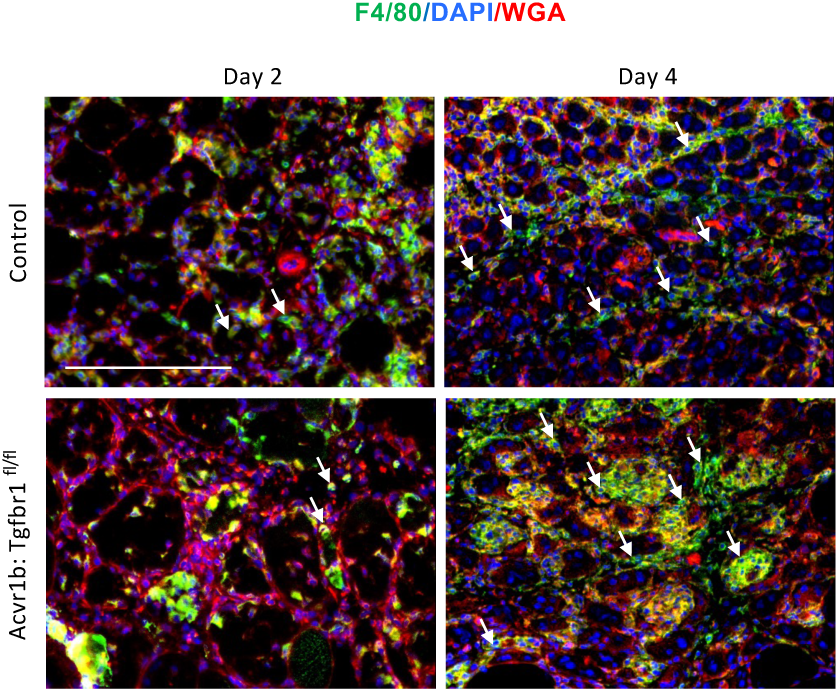
Number of macrophages in TA of Acvr1b^fl/fl^:Tgfbr1^fl/fl^ animals was increased 4 days post injury. Number of macrophages (F4/80^+^, green) in control and Acvr1b^fl/fl^:Tgfbr1^fl/fl^ animals on day 2 and 4 post injury. Nuclei were stained by DAPI (blue) and ECM of muscle were stained by WGA (red). Scale bar=100 μm.

**Figure 5-figure supplement 2.**
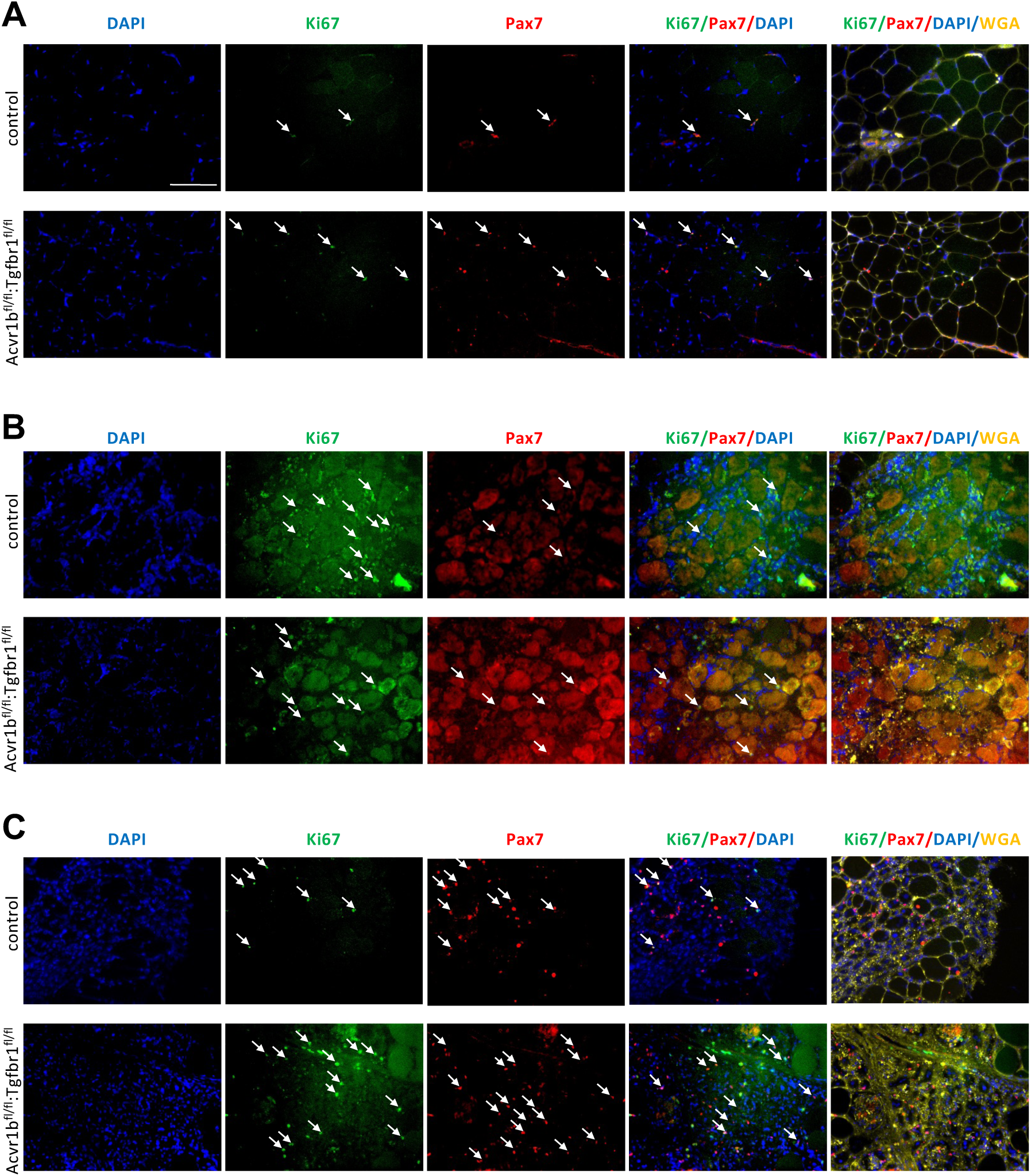
Increased number of proliferating cells and satellite cells in TA of Acvr1b^fl/fl^: Tgfbr1^fl/fl^ animals in absence of injury and 4 days post injury. (A) In absence of injury, proliferating cells (Ki67^+^, green, white arrows) and muscle satellite cells (Pax7^+^, red, white arrows) were shown by IF staining in low oxidative area of TA, where regeneration patches were found. Two (B) and 4 (C) days post injury, more Ki67^+^ and Pax7^+^ cells (white arrows) infiltrated in injured area. Nuclei were stained by DAPI (blue) and ECM of muscle were stained by WGA (yellow). Scale bar=100 μm. N=5-7.

**Figure 5-figure supplement 3.**
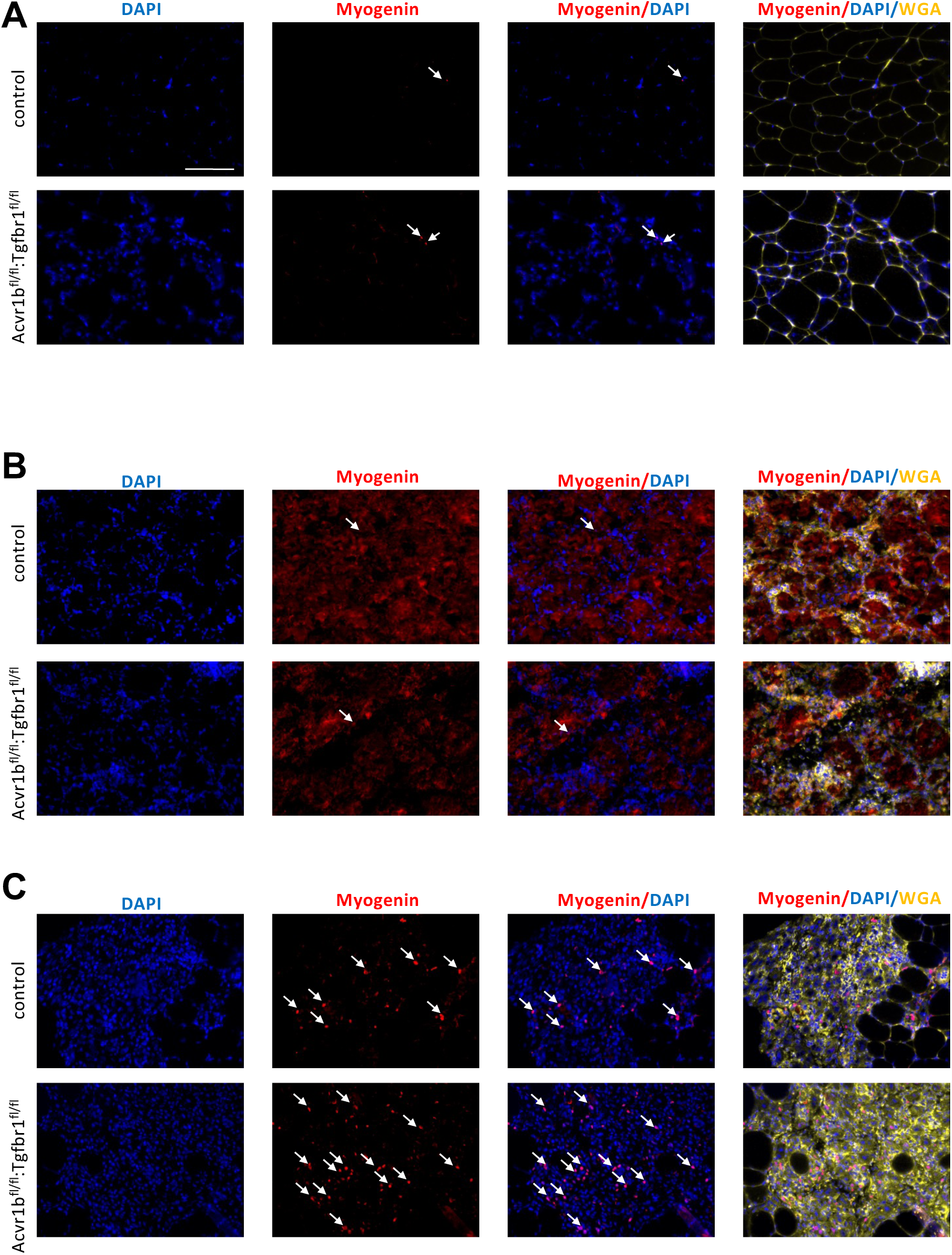
Increased number of differentiating muscle cells in TA of Acvr1b^fl/fl^: Tgfbr1^fl/fl^ animals 4 days post injury. On day 0 (A) and 2 days (B) post injury, number of differentiating myoblasts (myogenin^+^, red, white arrows) was of no difference between groups. Four days (C) post injury, more Myogenin^+^ cells were found in injured area of Acvr1b^fl/fl^: Tgfbr1^fl/fl^ animals. Nuclei were stained by DAPI (blue) and ECM of muscle were stained by WGA (yellow). Scale bar=100 μm. N=5-7.

